# Deciphering lentiviral Vpr/Vpx determinants required for HUSH and SAMHD1 antagonism highlights the molecular plasticity of these evolutionary conflicts

**DOI:** 10.1101/2024.03.07.583867

**Authors:** Pauline Larrous, Cassandre Garnier, Marina Morel, Michael M. Martin, Karima Zarrouk, Sarah Maesen, Roy Matkovic, Andrea Cimarelli, Lucie Etienne, Florence Margottin-Goguet

## Abstract

SAMHD1 and the HUSH complex constitute two blocks during primate lentivirus infection, the first by limiting reverse transcription and the second by inhibiting proviral expression. Vpr and Vpx of specific lentiviral lineages have evolved to antagonize these antiviral proteins. While the antagonism of SAMHD1 has been well characterized, the evolutionary and molecular determinants of the antagonism against HUSH are unknown. We used chimeric Vpr proteins between SIVagm.Ver and SIVagm.Gri lentiviruses infecting two African green monkey species, to investigate viral determinants involved in HUSH and SAMHD1 antagonisms. We found that different interfaces of closely related Vpr proteins are engaged to degrade different SAMHD1 haplotypes. In addition, we identified distinct viral determinants in SIVagm.Ver Vpr for SAMHD1 and HUSH degradation. The substitution of one residue in SIVagm.Gri Vpr is sufficient to gain the capacity to degrade SAMHD1, while the substitution of α-helix-3 confers HUSH antagonism. We also found that Vpx proteins of HIV-2 from people living with HIV have different abilities to degrade HUSH. These phenotypes rely on small changes in either the N or C terminal part of Vpx, depending on the context. On the host side, we found that HIV-2 and SIVsmm Vpx degrading HUSH from human and vervet monkey cells cannot not degrade HUSH in owl monkey cells, suggesting some host species-specificity.

Altogether, we highlight the molecular plasticity and constraints of viral proteins to adapt to host restrictions. HUSH, like SAMHD1, may have been engaged in ancient and more recent coevolution with lentiviruses and a player in viral fitness.

**IMPORTANCE:** Antiviral host proteins, the so-called restriction factors, block lentiviruses at different steps of their viral life cycle. In return, primate lentiviruses may counteract these immune proteins to efficiently spread *in vivo*. HIV-2 and some SIVs, but not HIV-1, inactivate SAMHD1 and HUSH, two host antiviral proteins, thanks to their Vpx or Vpr viral proteins. First, we uncovered here viral determinants involved in the function of closely related Vpr proteins from SIVs of African green monkeys and of HIV-2 Vpx alleles from people living with HIV-2. We show how these small viral proteins differently adapted to SAMHD1 polymorphism or to HUSH restriction and highlight their molecular plasticity. Finally, the capacity of divergent lentiviral proteins, including HIV-2 Vpx, to induce the degradation of HUSH depends of the cell/host species. Altogether, our results suggest that HUSH has been engaged in a molecular arms-race along evolution, and therefore is a key player in host-pathogens interaction.

## INTRODUCTION

Host restriction factors are antiviral proteins from the cell autonomous immunity that have been engaged in an evolutionary arms-race with the pathogenic viruses they have been fighting for millions of years (1, 2). They further represent molecular barriers to cross-species transmission of viruses (3–6). When transmitted to humans, the lentiviruses SIVcpz and SIVgor (from chimpanzees and gorillas, respectively) and SIVsmm (from sooty mangabeys) gave rise to HIV-1 and HIV-2, respectively. The different lentiviral lineages share a similar genomic organization, but differ in their set of accessory genes, which produce proteins largely dedicated to the counteraction of restriction factors and which have strongly evolved during lentiviral cross-species transmissions (6, 7). Determining the exact molecular residues/interfaces underlying these conflicts is therefore a major objective to better understand HIV-cell interactions and determinants of virus spillover.

All extant primate lentiviruses, including SIVagm (infecting African green monkeys), encode Vpr, which induces host G2/M cell cycle arrest (8–10). However, despite its importance in the dissemination and pathogenesis of SIVsmm (11), Vpx is found in only two of the eight major lineages of primate lentiviruses, HIV-2/SIVmac/SIVsmm (infecting humans, macaques and sooty mangabeys) and SIVrcm/mnd2 (infecting red-capped mangabeys and mandrills). *Vpr* and *vpx* genes are the results of duplication and recombination events of a precursor gene (reviewed in (12)). The encoded proteins share similarities in size (about 100 amino acids), structure (a N-terminal tail, 3 α-helices and a C-terminal tail) and functions. Nonetheless, they also present highly variable regions: VR1 upstream of helix 1, VR2 across the end of helix 2 and the beginning of helix 3, and VR3, which overlaps the C-terminal tail. In viral lineages that encode both Vpr and Vpx, Vpx induces the proteasomal degradation of the host restriction factor SAMHD1 (SAM and HD domain-containing protein 1), while in some other lineages that do not encode Vpx, such as SIVagm, the Vpr protein performs this function (13–15). Associated phylogenetic analyses showed that an ancestral Vpr protein acquired the anti-SAMHD1 activity prior to the molecular events that gave birth to Vpx (15). SAMHD1 is a 626 amino acid dNTPase that blocks viral DNA synthesis by lowering the pool of nucleotides in macrophages and quiescent CD4+ T cells (16, 17). By degrading SAMHD1, Vpx/Vpr proteins enable the virus to bypass a reverse transcription block. In this process, Vpx/Vpr directly binds SAMHD1 and bridges SAMHD1 to the DCAF1 adaptor of a Cullin4A-based ubiquitin ligase (13, 14, 18–21).

How Vpx interacts with SAMHD1 in a virus-host species-specific manner has been extensively studied. Strikingly, search for host determinants revealed that HIV-2/SIVsmm Vpx targets the C-terminus of SAMHD1, while SIVmnd2 and SIVrcm Vpx recognize its N-terminus (22–24). The divergence in SAMHD1 recognition is further witnessed by the presence of sites under positive selection during primate evolution in both N-and C-terminal domains of the protein (15, 22, 25). These site-specific adaptations in SAMHD1 are the results of host escape from viral antagonism, characteristic of a molecular virus-host arms-race.

The resolution of crystal structures and functional studies have allowed the identification of the interfaces between Vpx from different lineages and SAMHD1 (23, 24). In particular, a cluster of residues in the Vpx α-helix 2 of SIVmnd2 are in contact with SAMHD1, while these amino-acids are not involved in the case of Vpx from SIVsmm (23, 24).

By studying the coevolution between African green monkeys (AGMs) and their SIVs, Spragg and Emerman showed that SAMHD1 antagonism is crucial for viral fitness (26). AGMs comprise at least four closely-related species: *Chlorocebus tantalus*, *sabaeus*, *aethiops* (Grivet) and *pygerythrus* (Vervet). Each lineage of SIVagm, responsible of the natural infection of each species, has evolved to antagonize distinct SAMHD1 haplotypes through the use of Vpr (26). More precisely, among the seven SAMHD1 haplotypes identified in the AGMs, haplotype IV, but not V, is degraded by SIVagm.Ver Vpr, while the opposite is found for SIVagm.Gri Vpr; haplotype III is resistant to both Vprs, but is sensitive to SIVagm.Sab Vpr (26).

In addition to SAMHD1, HIV-2/SIVsmm Vpx and the Vpr from specific lineages can induce the degradation of the human HUSH complex (27, 28). The HUSH complex is composed of TASOR, MPP8 and periphilin and contributes to the silencing of cellular genes and retroelements with the help of MORC2(29). Due to HUSH degradation, Vpx/Vpr proteins favor viral expression in a model of HIV-1 latency (27, 28). Functional and evolutionary studies led us to conclude that HUSH antagonism is likely an ancient function of primate lentiviruses that preceded the birth of Vpx and SAMHD1 antagonism (15). In addition, HUSH antagonism appeared lentiviral species-specific, with only some Vpx/Vpr proteins degrading human HUSH. Whether lentiviral species-specificity is accompanied by host species-specificity, in line with host-virus competition along evolution, has not been investigated yet in the case of HUSH.

Here, we took advantage of the lentiviral-host specificity within the AGM lineage, both for SAMHD1 and HUSH antagonism, to identify viral determinants at stake. We found that the closely related Vpr proteins use different viral determinants to degrade different SAMHD1 haplotypes, highlighting the molecular plasticity and adaptation of the virus to the host. In addition, viral determinants against HUSH are different from those against SAMHD1. We further studied Vpx proteins of HIV-2 from people living with HIV-2 (PLWH-2), which were competent for SAMHD1 degradation (30). We found that some Vpx proteins can induce HUSH degradation, while others cannot, and this diversity seems independent of the corresponding status of the viremia. Depending on the Vpx, restoration of the capacity to induce HUSH degradation was obtained by changing either the N-terminal part or the C-terminal part of the viral protein.

Finally, we describe one evidence of host-species specificity with HIV-2/SIVsmm Vpx unable to counteract HUSH in New World monkey cells. Altogether, our results suggest the existence of a dynamic interplay between HUSH, SAMHD1 and primate lentiviruses along evolution.

## MATERIALS AND METHODS

### Plasmids

Vpr SIVagm.ver9063 (KF741096), Vpr SIVagm.gri677 (sequence as the one used in (26)), Vpr SIVagm.Tan1 (U58991) and Vpr SIVagm.Sab1 (S46351) chimera 1 to 10 (sup Fig 1), Vpx proteins of HIV-2 from PLWH-2 and Vpx chimeras have been synthesized after codon-optimisation and subcloned into the pAS1b vector (pAS1b-HA) to get HA-epitope-tagged proteins (HA at the N-terminus). Vpr SIVagm.Tan1 is also expressed in pCDNA3-3xFlag vector with a Flag epitope at the N-terminus. The mutants of Vpr were produced by site-directed mutagenesis according to Phusion polymerase manufacture guide (Thermofisher) or CloneAmp HiFi polymerase manufacture guide (Takkara) using Vpr SIVagm.ver9063 or chimera 8 or 9 in the pAS1B vector as templates. Lentiviral proteins Vpx HIV-2 Gh (P18045.1), Vpx SIVsmm (P19508.1), Vpx SIVmnd2.GAx14 (AAK82846.1), Vpx SIVrcm.NG411 (AAK69676.1) and Vpx SIVrcm.Gab1 (AAM34564.1) are also expressed from the pAS1b vector (HA tag at the N-terminus). Human TASOR (NP_001106207.1) and Owl monkey TASOR (NCBI References Sequences: XP_012316204) are expressed from vectors pLenti-Flag and pCMV6-Flag respectively, in fusion with the myc-DDK epitope at the C-terminus. Constructs expressing Haplotypes III, IV and V of SAMHD1 from AGMs are gifts from M. Emerman and are expressed with a HA epitope at the C-terminus from the pLPCX vector (KF741043, KF741044 and KF741045).

**Figure 1:**
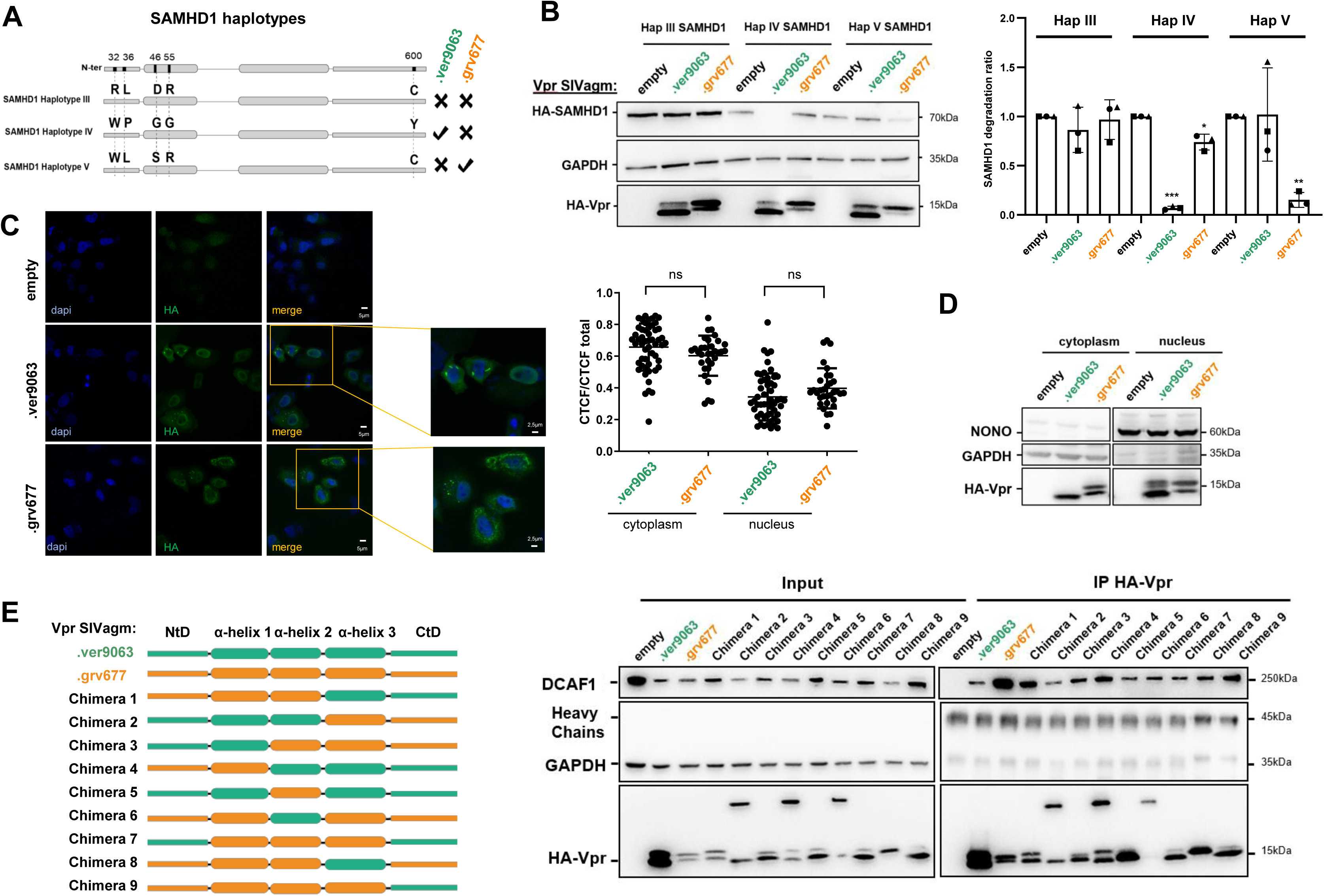
Closely related Vpr proteins, which differently antagonize SAMHD1 variants, bind DCAF1. (A) Schematic representation of SAMHD1 haplotypes III, IV and V. The phenotype of their degradation as shown in B is indicated on the right (v: degradation, x: no degradation). (B) 293T cells stably expressing HA-tagged SAMHD1 haplotypes III, IV and V were transfected with plasmids encoding SIVagm Vpr proteins (ver9063: strain of SIVagm.Ver, grv677: strain of SIVagm.Gri). (Left) The indicated proteins were revealed by western-blot (one representative western-blot is shown). (Right) Quantification of SAMHD1 degradation rate from 3 experiments. A one-sample t-test was performed. Data are presented as mean ± standard deviation (SD). Statistical significance is indicated as follows: *p < 0.05, **p < 0.01, ***p < 0.001. (C) HA-Vpr proteins were co-expressed in HeLa cells cultured on glass slides. Cells were fixed and stained with fluorescent probes for DNA (DAPI; blue), HA-Vpr (anti-HA Alexa Fluor 488-conjugated; green). (Left) Images were acquired using confocal microscopy, representative images showing the distribution of HA-Vpr are shown. (Right) Quantification of the mean fluorescence intensity per cell (n = 30 cells) using ImageJ. Error bars represent the standard deviation. No statistically significant difference in fluorescence intensity was observed between SIVagm.Ver, and SIVagm.Gri Vpr (p > 0.05, Student’s t-test). (D) HeLa cells were transfected with plasmids encoding SIVagm Vpr proteins. Cells were fractionated before the lysis of the nucleus and the cytoplasm. The indicated proteins were revealed by western blot. (E) Schematic representation of SIVagm Vpr proteins and chimeras. The scale is respected for the length of the different domains. (F) Indicated HA-Vpr constructs were expressed in HeLa cells, then an anti-HA immunoprecipitation was performed, and proteins were revealed by western-blot.

### Cell culture

Cells were tested regularly for mycoplasma contaminations; experiments were only performed on non-contaminated cells. ATCC-purchased HeLa (CCL-2), VERO (CCL-81), HEK293T (CLR-3216), HEK293FT (293T cells optimized for VLP production, gift from N. Manel) and OMK cells were cultivated in media DMEM (Thermofisher) containing 10% fetal bovine serum (FBS, Eurobio), 1,000 units/mL penicillin, 1,000µg/mL streptomycin (Life Technologies). J-Lat A1 (gift from E. Verdin) were cultivated in media RPMI (Thermofisher) supplemented as described for the DMEM medium and, in addition, 2mM glutamine (Life Technologies). HEK293T stable cell lines expressing the haplotypes III, IV and V of AGM SAMHD1 were generated by transduction of VLP containing pLPCX-Hap III, pLPCX-Hap IV and pLPCX-Hap V and cultivated four days before puromycin selection.

### siRNA treatment

siRNA transfections were performed with DharmaFECT1 (Dharmacon, GE Lifesciences). The final concentration for all the siRNA was 40nM. The following siRNA were purchased: siTASOR: SASI_Hs02_00325516 (Sigma Aldrich); siDCAF1: J-021119-10-0005 (Dharmacon) and the non-targeting control siRNA: MISSION siRNA Universal Negative Control 1, SIC001 (Sigma Aldrich).

### Virus-Like-Particle production, delivery and transduction

VLPs were produced in HEK293FT cells by co-transfection by the calcium-phosphate co-precipitation method of VSV-G plasmid (3µg), SIV3+ ΔVpr ΔVpx vectors (8µg) and 8µg of pAS1b-HA-Vpr (or chimeric proteins or point mutants) or pAS1b-HA-Vpx or pAS1b-HA (empty) and in some experiments with a transfer gene pGAE1.0 (SIVmac-CMV-GFP) used as a reporter gene. SIV3+ ΔVpr ΔVpx packaging vector is a gift from N. Landau and is described in Gramberg *et al* (31). VLPs used for the establishment of the HEK293T stable cell lines expressing HA-SAMHD1 (agm) were produced with VSV-G plasmid (3µg), pHIT60 MLV packaging vector (8µg) and 8µg of pLPCX vector expressing Hap III or Hap IV or Hap V. In both cases, 3.10^6^ cells were plated in 10cm culture dishes the day prior transfection. Cell culture medium was collected 48h after transfection and filtered through 0,45µm pores filters. For SAMHD1 VLP, 10mM NaBu were added 24h after transfection and the cells washed at the end of the day. VLP were concentrated by sucrose gradient and ultracentrifugation (1h30 at 100,000g). The incorporation of the viral proteins was assessed by western blot and a quantification of the level of HIV-2 capsid (P27) and HA-Vpr was performed to deliver the same quantity of viral proteins. J-Lat A1 cells were treated with VLP for 7h in reduced medium prior to overnight TNFα (1ng/mL) treatment. VERO and OMK cells were plated in 12-well dishes at a density of 3.10^5^ cells and transduced the day after in reduced medium, cells were harvested the day after transduction.

### Flow cytometric analyses

J-Lat A1 cells were collected and resuspended in PBS-EDTA (0,5mM). Data were collected and analyzed with BD Accuri C6 cytometer or Attune and software CFlow Plus or FlowJo. At least 10,000 events in P1 were collected, the GFP-positive population was determined using untreated J-Lat A1 cells as their level of GFP expression is low. The same gate was maintained for all conditions and analysis were performed overall GFP-positive population.

### SAMHD1 degradation assay

HEK293T Hap III, Hap IV and Hap V were plated in 12-well dishes at 1,5.10^5^ cells and transfected the following day using the calcium-phosphate co-precipitation method. Different amounts of the pAS1B vector expressing the different Vpr proteins (1 to 2µg) were transfected to get the same level of expression of the viral proteins (with adjustment to get the same level of total DNA per condition). Cells were harvested 48h post-transfection for western blot analysis. Lysis of the cells was performed in 100µL of RIPAC buffer (50mM Tris-HCl pH7.5, 150mM NaCl, 10% Glycerol, 2mM EDTA, 0.5% NP40, 0.1% SDS) containing an anti-protease cocktail (A32965, Thermofisher), lysates were centrifuged at 16,000g for 10min to remove cell debris.

### Cell fractionation

All immunoprecipitation experiments are performed in the nuclear fraction of HeLa cells. Cells grown in 10cm dishes were washed with cold Dulbecco’s PBS 1x (ThermoFisher). After trypsinization (Thermofisher), cells were recovered in 1,5mL tubes and washed once with ice-cold PBS. After 4 min of centrifugation at 400xg, 1mL of cytoplasmic lysis buffer (10 mM TRIS-HCl pH7.5, 10 mM NaCl, 3 mM MgCl2, and 0.5% IGEPAL® CA-630 (I8896-100ML Merck)) was added on the cell pellet and resuspended pellet was incubated on ice for 5 min. Cells were then centrifuged at 300g for 4 min at 4 °C and the supernatant was saved for cytoplasmic fraction. The pellet was washed with 1mL of cytoplasmic lysis buffer and re-centrifuged at 300g for 4 min at 4 °C. Finally, the nuclear pellet was lysed with 300μL of RIPA buffer.

### Immunoprecipitation assay, western blot procedure and antibodies

For HA-Vpr (AGM WT or chimeric proteins) and TASOR-Flag immunoprecipitations, HeLa cells were plated at respectively 2,5.10^6^ cells in 10cm dishes and co-transfected by the calcium-phosphate co-precipitation method with pAS1b-HA or pAS1b-HA-Vpx or Vpr (SIVagm WT or chimeric proteins) and plenti-TASOR-FLAG or pCMV6-TASOR-FLAG. Cells were harvested 48h post-transfection for western blot analysis. Lysis of the cells was performed in 700µL of RIPA buffer (50mM Tris-HCl pH7.5, 150mM NaCl, 10% Glycerol, 2mM EDTA, 0.5% NP40) containing an anti-protease cocktail (A32965, Thermofisher) and spun at 16,000g for 10min to remove cell debris. 500µg of cell lysates were incubated with pre-washed EZview Red ANTI-HA or FlagM2 affinity Gel Beads (E6779 and F2426, Merck) at 4 °C under overnight rotation. After three washes in wash buffer (50mM Tris-HCl pH7.5, 150mM NaCl), immunocomplexes were eluted with Laemmli buffer 1X with 20mM DTT and were separated by SDS-PAGE (Bolt Bis-Tris, 4-12%, Life Technologies). Following transfer onto PVDF membranes, proteins were revealed by immunoblot and signals were acquired with Fusion FX (Vilber Lourmat). The following antibodies, with their respective dilution in 5% skimmed milk in PBS-tween 0.1% were used: anti-HA-HRP (3F10) (N°12013819001, Roche) 1/10,000; anti-FLAG-HRP (A-8592, lot 61K9220, Sigma) 1/10,000; anti-HA (HA-7, H3663, lot 066M4837V, Merck) 1/1,000; anti-Flag M2 (F1804-200UG-lot SLCD3990, Merck) 1/1,000; anti-TASOR (HPA006735, lots A106822, C119001, Merck) 1/1,000; anti-DCAF1 (11612-1-AP, ProteinTech) 1/1000; Anti-p27/p55 and anti-P24 were provided by the NIH AIDS research and reference reagent program (ref ARP392/393) 1/1000; anti-βActin (AC40, A3853, Merck) 1/1000; anti-GAPDH (6C5, SC-32233, Santa Cruz) 1/1,000. All HRP-conjugated secondary antibodies, anti-mouse (31430, lot VF297958, Thermofisher) and anti-rabbit (31460, lots VC297287, UK293475 Thermofisher), were used at a 1/20,000 dilution before reaction with Immobilon Classico (WBLUC0500, Merck Millipore) or Forte (WBLUF0100, Merck Millipore) Western HRP.

### Immunofluorescence assay

HeLa cells were cultivated on glass side and transfected as explained before. Cells were fixed with 4% paraformaldehyde for 15 min and permeabilized with 0.1% Triton for 15 min. Blocking step was performed with 2% bovine serum albumin solution for 1h at room temperature. Cells were incubated with antibody against anti-hemagglutinin mouse IgG monoclonal conjugate Alexa Fluor 488 conjugated (Invitrogen) for 1h. The cells were washed and the nuclei counter stained with 40-6-diamidino-2-phenylindole (DAPI, Sigma-Aldrich) for 20 min. The coverslip cells were mounted with Prolong gold/diamond antifade reagent (Invitrogen). Immunofluorescence images were captured by using a Leica DMI6000 confocal microscope at the IMAG’IC core facility.

## RESULTS

To identify viral determinants involved in SAMHD1 and HUSH antagonism, we leveraged the lentiviral-host specificity within the SIV-AGM lineage. Among the seven SAMHD1 haplotypes identified in the AGM population (26), we chose to study three haplotypes (III, IV and V) that present no more than four amino acid differences and that have been described as: abundant in the Vervet but very rare in the Grivet AGM species (haplotype IV), or abundant in the Grivet and absent in the Vervet AGM species (haplotype V), or absent/rare in the Grivet/Vervet AGM species (haplotype III) (Fig. 1A). The ability of Vpr proteins from SIVagm viruses to degrade the different SAMHD1 haplotypes was then examined (Fig. 1A and 1B). To this end, 293T stable cell lines encoding the different HA-tagged SAMHD1 haplotypes were first established and Vpr was then expressed ectopically by transient DNA transfection (primary sequences of Vpr proteins in Sup Fig. 1). Of note, the sequence of SIVagm.Gri Vpr (SIVagm.grv677 Vpr) harbors two amino-acid changes (A2T and R102G) compared to the sequence of the original protein (NCBI RefSeq NP_054371.1), which were acquired after virus isolation (26, 32). The sequence of SIVagm.Ver Vpr is from SIVagm.Ver9063 (GenBank KF741096.1). Under these conditions, the Vpr protein derived from the SIVagm.Ver9063 was able to induce the degradation of SAMHD1 haplotype IV, but not haplotype V (Fig. 1B, quantification on the right). Conversely, the Vpr protein derived from the SIVagm.grv677 led to the degradation of SAMHD1 haplotype V but not haplotype IV (Fig. 1B). As previously shown by Spragg and Emerman (26), neither of the two Vpr proteins induced the degradation of SAMHD1 haplotype III, which is predominantly found in Sabaeus AGM species (Fig. 1B). The observed specificities of Vpr proteins could not be explained by major subcellular localisation differences, because both Vprs exhibited similar distribution in the nucleus and the cytoplasm, following immunofluorescence and biochemical fractionation experiments (Fig. 1C and 1D). However, the specificities could be the result of post-translational modifications, as at least two bands were revealed in different proportions for SIVagm.Ver Vpr and SIVagm.Gri Vpr, but this was not further investigated (Fig. 1B and 1D). Overall, phenotypes of SAMHD1 degradation suggest the existence of specific interactions between SAMHD1 haplotypes present in a given species and the Vpr protein from the SIVagm infecting the same species.

SIVagm.Ver and SIVagm.Gri Vpr proteins share 69% of identity at the amino acid level. To decipher the viral determinants involved in SAMHD1 antagonism, we constructed Vpr protein chimeras by exchanging their different α-helices, N-and C-termini domains (NtD and CtD) (Fig. 1E, Sup Fig. 1 for sequences). Some chimeras harbored a dimeric form in addition to the monomeric form, but all were able to bind DCAF1 (Fig. 1F). Chimera 6 was hardly present as a monomer, possibly impacting its activity (Fig. 1F). All chimeras able to induce SAMHD1 haplotype IV degradation in human cells contained the CtD of SIVagm.Ver Vpr (chimeras 1, 4, 5, 7) (Fig. 2A). The replacement of the CtD from SIVagm.Gri by the CtD from SIVagm.Ver (chimera 9) conferred the SIVagm.Gri Vpr the ability to induce SAMHD1 haplotype IV degradation (gain-of-function, Fig. 2A). These results suggest that the integrity of the C-terminal domain of SIVagm.Ver Vpr is a critical determinant for SAMHD1 haplotype IV degradation. Furthermore, the CtD of SIVagm.Tan Vpr was also able to confer the SIVagm.Gri protein (chimera 10) the ability to degrade SAMHD1 haplotype IV (Fig. 2B). Therefore, SIVagm Vpr proteins that target the C-terminus of SAMHD1 (SIVagm.Tan and SIVagm.Ver Vpr (26)) rely on C-terminal Vpr determinants to induce SAMHD1 haplotype IV degradation.

**Figure 2:**
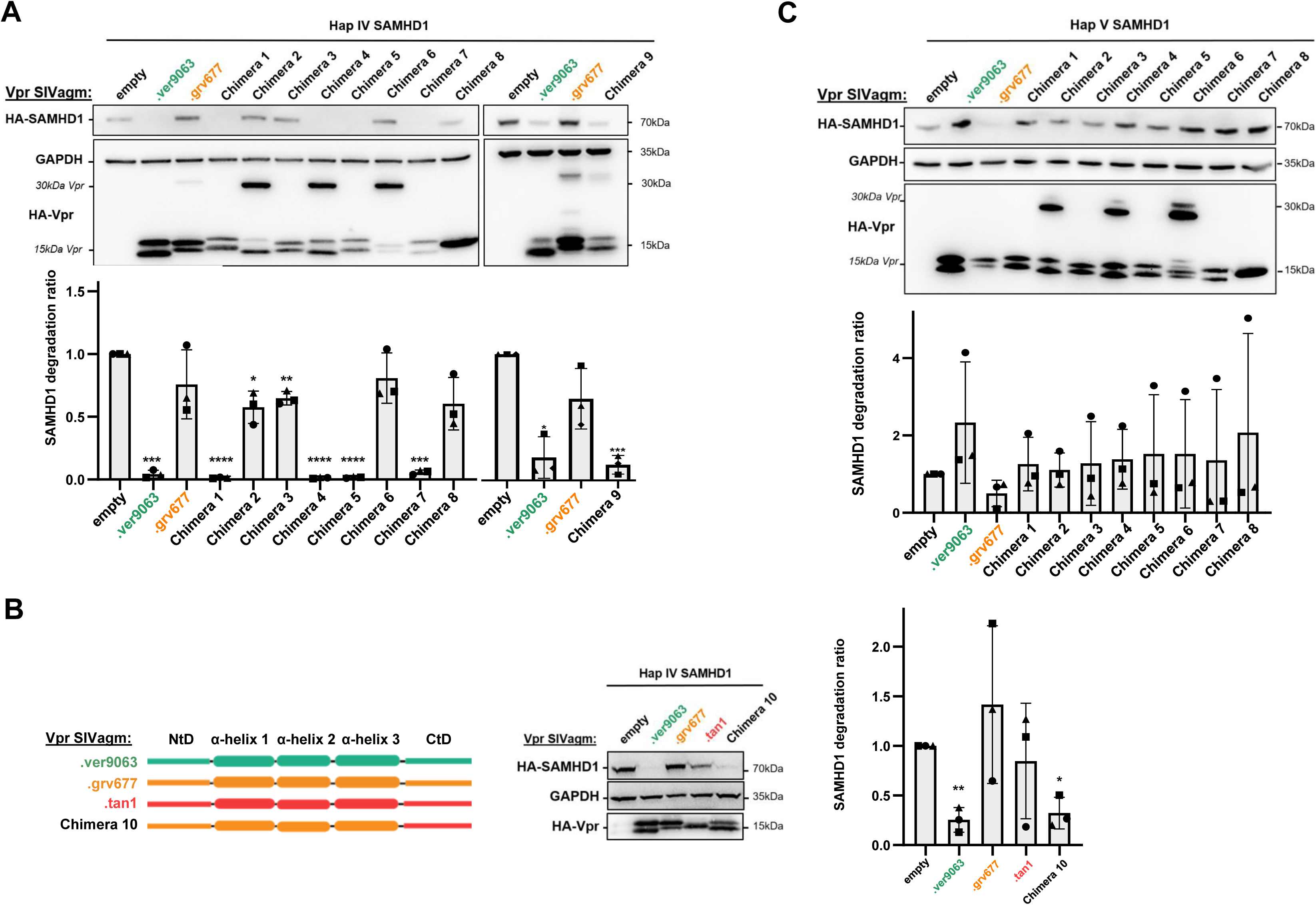
Closely related Vpr proteins can induce the degradation of SAMHD1 variants (Haplotypes IV and V) through distinct molecular determinants. (A) 293T cells stably expressing HA-tagged SAMHD1 haplotype IV were transfected with plasmids encoding SIVagm Vpr proteins and chimeras. (Top) A representative western blot from 3 experiments is shown, (bottom) Quantification of SAMHD1 degradation rate from 3 experiments. A one-sample t-test was performed. Data are presented as mean ± standard deviation (SD). Statistical significance is indicated as follows: *p < 0.05, **p < 0.01, ***p < 0.001. (B) Same as in (A) but with different Vpr constructs as indicated. (C) Same as in (A) but with cells stably expressing SAMHD1 haplotype V.

None of the viral chimeras led to consistent degradation of SAMHD1 haplotype V, suggesting that determinants throughout the viral protein, or the protein’s complete conformation, are essential for this activity (Fig. 2C). Despite all chimeras were able to bind DCAF1, we cannot exclude that the lack of degradation might result from a global structural defect. Therefore, only chimeras able to degrade either SAMHD1 haplotype IV (1,4,5,7,9,10) or TASOR (1,4,5,8, see below) were taken into account to draw conclusions. With this in mind, our results show that different viral interfaces in SIVagm.Ver Vpr and SIVagm.Gri Vpr are involved in the degradation of SAMHD1 haplotypes IV and V, respectively.

We then used the same Vpr chimeras to identify the viral determinants involved in the degradation of TASOR, the core component of the HUSH complex (33). First, we assayed the degradation of endogenous TASOR in AGM Vervet cells (VERO cells). Vpr proteins were incorporated into Viral-Like Particles (VLPs) and then delivered in VERO cells. Vpr incorporation into VLPs was quantified by Western-blot, and VLPs quantities were adjusted to deliver similar amounts of viral proteins into cells (Fig. 3A). SIVagm.Ver, SIVagm.Sab and SIVagm.Tan Vpr proteins were able to induce the degradation of endogenous TASOR in Vervet cells, in contrast to the SIVagm.Gri Vpr (Fig. 3B). Of note, our TASOR antibody detected two bands in VERO cells, but subsequent TASOR siRNA experiment suggested that only the lower band was indeed corresponding to TASOR (Fig. 3B, right). As previously described, similar phenotypes of TASOR degradation were found in human J-Lat A1 cells (Fig. 3C), a Jurkat T-cell line derivative that harbors a latent HIV-1 mini-genome expressing GFP under the control of the proviral LTR promoter (27, 34). In this HIV-1 latency model, TASOR degradation correlated with an increase of the percentage of GFP-positive cells, indicative of the reactivation of the latent provirus (Fig. 3D). Chimeras were further tested in both cell types, VERO and human cells (Fig. 4). All chimeras were well incorporated into VLPs, except chimera 2 (Fig. 4A). Degradation phenotypes were the same in Vero and J-Lat A1 cells: all the Vpr constructs that induced TASOR degradation contained the SIVagm.Ver Vpr α-helix 3 (chimeras 1, 4, 5 and 8) (Fig. 4B for Vero cells and 4C for human J-Lat A1 cells). In particular, chimera 8, which harbors the SIVagm.Ver α-helix 3 in a SIVagm.Gri Vpr background, was competent for TASOR degradation (gain-of-function), indicating that the SIVagm.Ver Vpr α-helix 3 region is a key viral determinant for TASOR degradation. As expected, TASOR degradation correlated with reactivation in the J-Lat-A1 model (Fig. 4D). Altogether, the C-terminal domain of SIVagm.Ver Vpr appears as a critical determinant for SAMHD1 haplotype IV degradation, while α-helix 3 is a key determinant for TASOR degradation.

**Figure 3:**
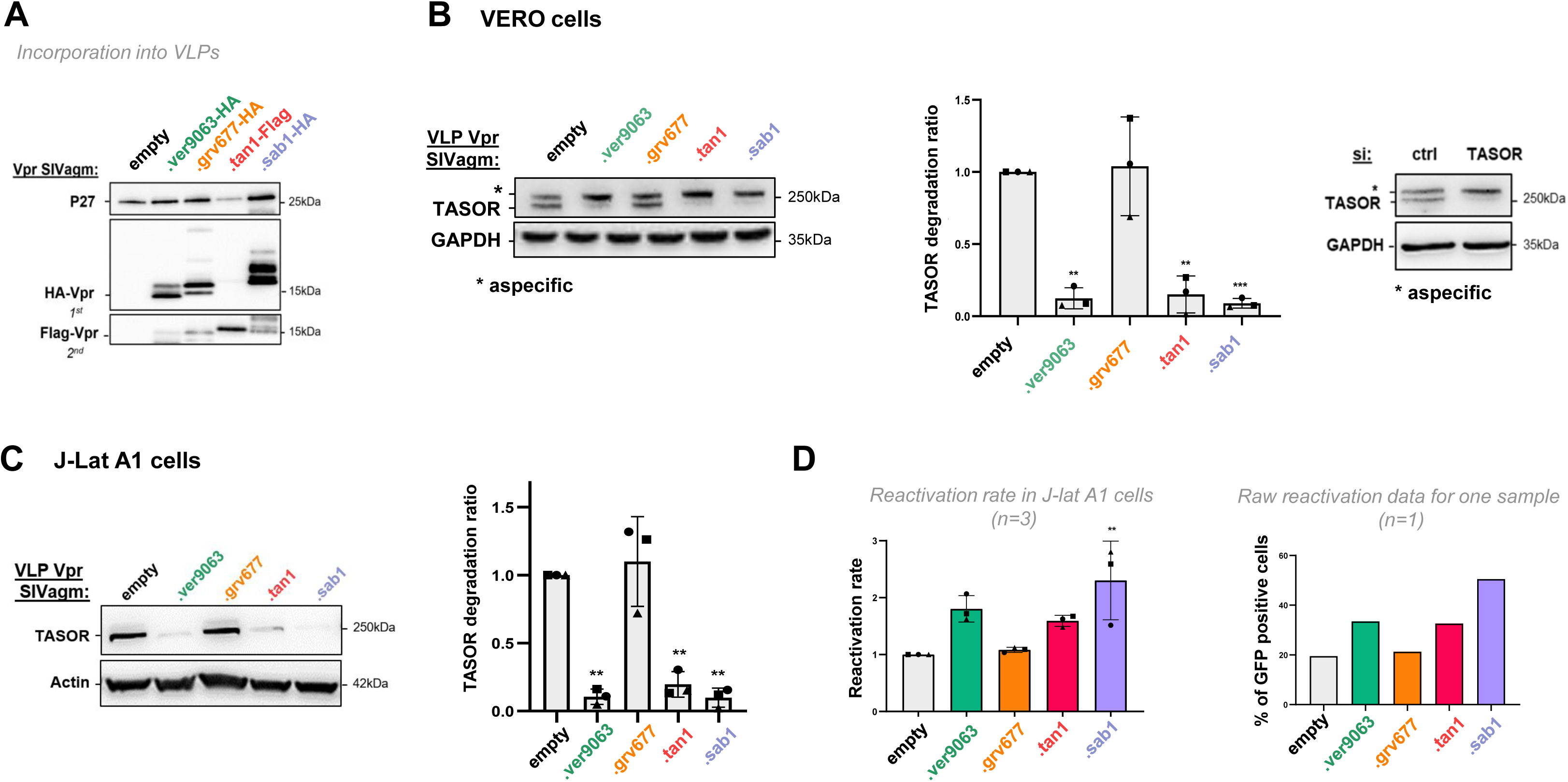
SIV strains of African green monkey species differently degrade TASOR both in Vero and human cells. (A) VLPs were produced in 293FT by co-transfection of a packaging vector, an envelope VSVg vector and a vector encoding HA-or Flag-Vpr as indicated. 48h post transfection, supernatants were harvested and VLPs, concentrated by ultracentrifugation, were analyzed by western blot. VLP production was checked with anti-P27 (capsid) antibody and HA-Vpr incorporation with an anti-HA antibody. (B) (Left) VERO cells were treated with VLPs containing SIVagm Vpr proteins as indicated and whole-cell extracts analyzed by western blot. (Middle) Quantification of TASOR degradation rate from 3 experiments. A one-sample t-test was performed. Data are presented as mean ± standard deviation (SD). Statistical significance is indicated as follows: *p < 0.05, **p < 0.01, ***p < 0.001. (Right) VERO cells were treated with either siRNA CTRL or siRNA TASOR. (C) and (D) Human J-Lat A1 T cells were treated with VLPs containing SIVagm Vpr proteins and stimulated overnight with TNF-α, then cells were analyzed by western-blot and flow cytometry for the percentage of GFP-positive cells. (C) (Left) Whole-cell extracts were analyzed by western blot, the immunoblot shown is representative of at least 3 independent VLP productions. (Right) Quantification of TASOR degradation rate from 3 experiments. A one-sample t-test was performed. Data are presented as mean ± standard deviation (SD). Statistical significance is indicated as follows: *p < 0.05, **p < 0.01, ***p < 0.001. (D) The reactivation rate corresponds to the percentage of GFP-positive cells in the presence of one viral protein over the percentage obtained without viral protein (empty condition). Reactivation rates from 3 independent experiments are shown, along with raw reactivation data from one representative experiment. A one-sample t-test was performed, with statistical significance denoted as: *p < 0.05, **p < 0.01, ***p < 0.001.

**Figure 4:**
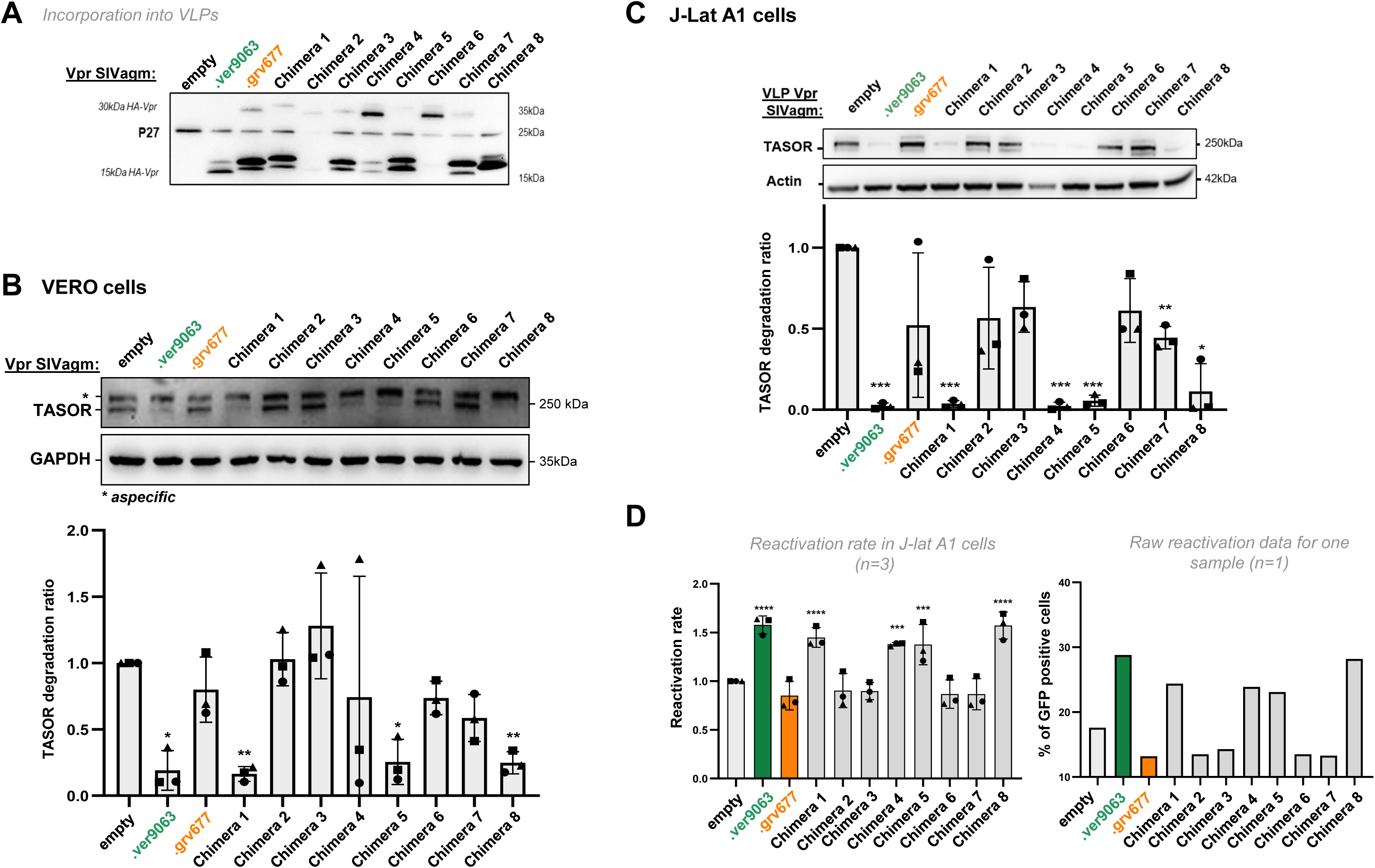
α-helix 3 of SIVagm.Ver Vpr confers on the SIVagm.Gri protein the ability to induce the degradation of the HUSH core protein TASOR. (A) Incorporation of Vpr chimera proteins into VLPs as described in Figure 3A. (B) VERO cells were treated with VLPs containing SIVagm Vpr proteins. After overnight treatment cells were lysed. (Top) Whole-cell extracts were analyzed by western blot, the immunoblot shown is representative of at least 3 independent VLP productions. (Bottom) Quantification of TASOR degradation rate from 3 experiments. A one-sample t-test was performed. Data are presented as mean ± standard deviation (SD). Statistical significance is indicated as follows: *p < 0.05, **p < 0.01, ***p < 0.001. (C) Human J-Lat A1 T cells were treated with VLPs containing SIVagm Vpr proteins. After overnight treatment with TNF-α, cells were analyzed by flow cytometry for the percentage of GFP-positive cells. (Top) Whole-cell extracts were analyzed by western blot, the immunoblot shown is representative of at least 3 independent VLP productions. (Bottom) Quantification of TASOR degradation rate from 3 experiments. A one-sample t-test was performed. Data are presented as mean ± standard deviation (SD). Statistical significance is indicated as follows: *p < 0.05, **p < 0.01, ***p < 0.001. (D) The reactivation rate corresponds to the percentage of GFP-positive cells in the presence of one viral protein over the percentage obtained without viral protein (empty condition). Reactivation rates from 3 independent experiments are shown (left), along with raw reactivation data from one representative experiment (right). A one-sample t-test was performed, with statistical significance denoted as: *p < 0.05, **p < 0.01, ***p < 0.001.

To further delineate key residues, we performed sequence analyses of the viral proteins in the two regions. We first searched for potential “loss of function” mutations, which would impair SAMHD1 haplotype IV degradation, in the few residues that differ between the CtD of SIVagm.Ver and SIVagm.Gri Vprs proteins (Fig. 5A). We made the corresponding Vpr mutants by changing residues in the CtD of chimera 9, thus producing chimera 9 Q94G, E97S and RANRA-APPP (with RANRA residues at position 110 changed to APPP). We also made the R102G mutant directly in the SIVagm.Ver Vpr. Of note, D104 and D119 were not changed, because they are both present in the phenotypically different Vprs. The resulting proteins were all able to induce SAMHD1 haplotype IV degradation, except SIVagm.Ver Vpr R102G (Fig. 5B). Furthermore, the reciprocal G102R change in SIVagm.Gri Vpr restored its ability to induce SAMHD1 haplotype IV degradation, highlighting R102 as a key residue for this activity (Fig. 5C). Of note, the G102R substitution allowed to recover the original sequence of SIVagm.Gri Vpr; the original protein is now notified with an asterisk throughout the manuscript (i.e., SIVagm.grv677* Vpr). Results are summarized in Figure 5D.

**Figure 5:**
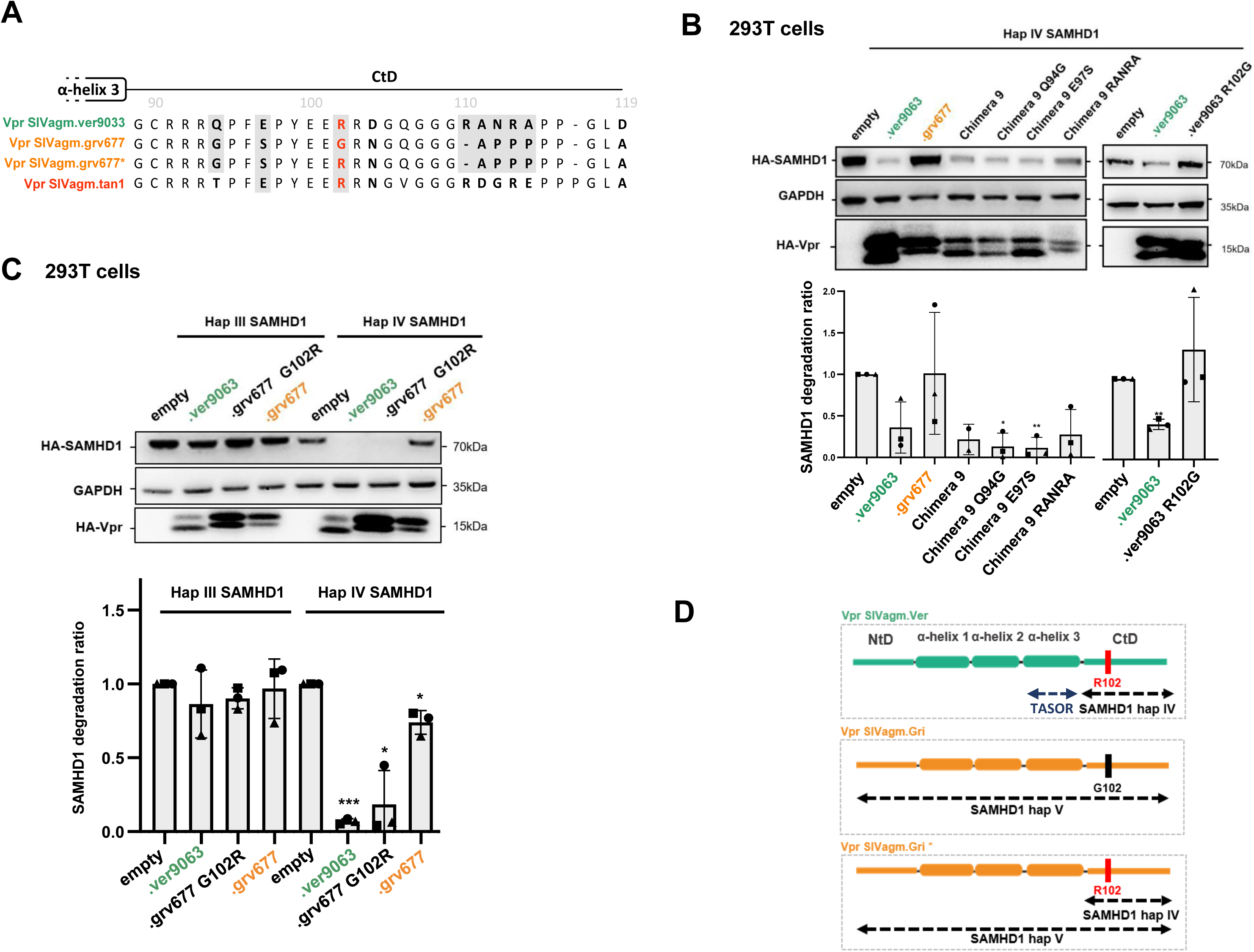
The substitution of only one amino-acid in SIVagm.Gri Vpr restores its ability to induce the degradation of SAMHD1. (A) The C-terminal domain sequences from SIVagm.Ver9063, SIVagm.Grv677, SIVagm.Grv677* (original sequence) and SIVagm.Tan Vpr proteins were aligned to point out amino acid differences. Substitutions tested in SIVagm.Ver Vpr to assess a functional loss are highlighted in grey. The key residue identified for haplotype IV SAMHD1 degradation is shown in red. (B) 293T cells were transfected with plasmids coding for HA-tagged Vpr proteins, chimera 9 or mutants, as indicated; (top) extracts from these cells were analyzed by western blot, one representative experiment is shown. (bottom) Quantification of SAMHD1 degradation rate from 3 experiments. A one-sample t-test was performed. Data are presented as mean ± standard deviation (SD). Statistical significance is indicated as follows: *p < 0.05, **p < 0.01, ***p < 0.001. (C) SIVagm.Ver Vpr, SIVagm.Gri Vpr and SIVagm.Gri Vpr G102R were overexpressed by transfection in 293T cells stably expressing the haplotype III or IV of HA-tagged SAMHD1. (top) Extracts from these cells were analyzed by western blot, one representative experiment is shown. (bottom) Quantification of SAMHD1 degradation rate from 3 experiments. A one-sample t-test was performed. Data are presented as mean ± standard deviation (SD). Statistical significance is indicated as follows: *p < 0.05, **p < 0.01, ***p < 0.001. (D) Schematic representation of key viral determinants involved in SAMHD1 and TASOR degradation.

Similarly, we searched for potential “loss of function” mutations, which would impair TASOR degradation, by analyzing differences in α-helix 3 (Fig. 6A). Six residues that were different between SIVagm.Ver and SIVagm.Gri Vpr proteins in the α-helix 3 were changed two by two. Incorporation of the corresponding viral proteins into VLPs was checked (Fig. 6B). All the corresponding mutants in the SIVagm.Ver Vpr background retained the capacity to degrade human TASOR and to reactivate HIV-1 in the J-Lat A1 model (Fig. 6C and 6D). Because several residues in α-helix 3 of Vpr/Vpx proteins are required for DCAF1 binding (19, 23, 24), we wondered whether the defect in TASOR degradation could result from a defect in DCAF1 binding. However, we found that SIVagm.Ver and SIVagm.Gri Vpr proteins both bound DCAF1 suggesting that differences in TASOR degradation are not linked to DCAF1 binding (Fig. 1F or 6E). Interestingly, exogenously-expressed Flag-tagged human TASOR seemed to better interact with SIVagm.Ver Vpr than with SIVagm.Gri Vpr (Fig. 6E). Vpr differences in TASOR degradation could then result from differences in TASOR binding. Mutations two by two in α-helix 3 did not impair DCAF1 or TASOR-Flag binding in agreement with degradation assays (Fig. 6E). Thus, α-helix 3 appears essential for TASOR degradation, but we could not further narrow-down the specific residues.

**Figure 6:**
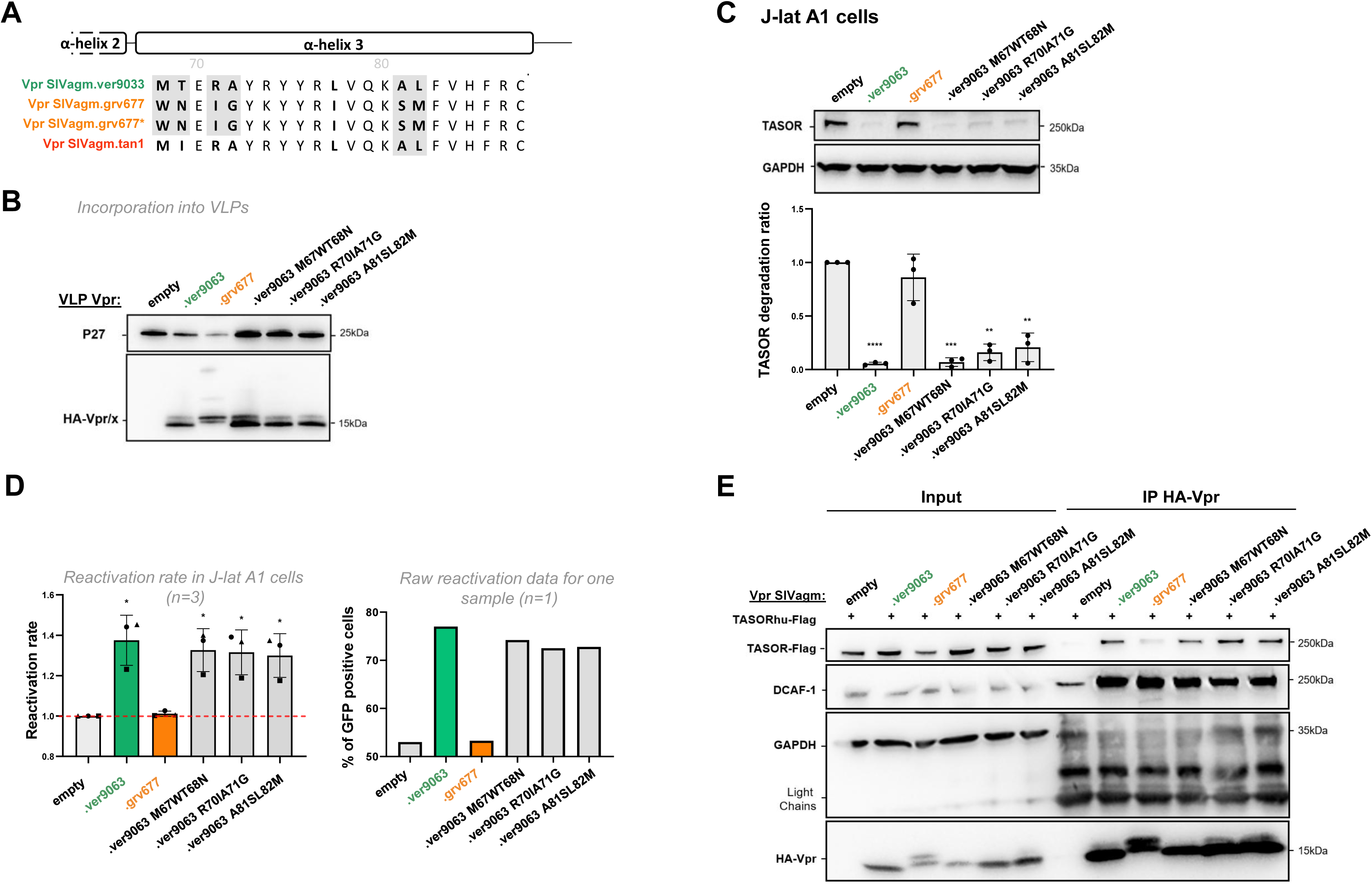
The whole integrity of SIVagm.Ver Vpr α-helix 3 contributes to the degradation of TASOR. (A) α-helix 3 sequences from SIVagm.Ver9063, SIVagm.Grv677, SIVagm.Grv677* (original sequence) and SIVagm.Tan Vpr proteins were aligned to point out amino acid differences. Substitutions tested in SIVagm.Ver to assess a functional loss are highlighted in grey. Key residues identified for TASOR degradation are shown in red. (B) Same as Fig. 3A with SIVagm.Ver9063 mutants. (C) Human J-Lat A1 T cells were treated with VLPs containing Vpr AGM proteins. After overnight treatment with TNF-α, (top) whole-cell extracts were analyzed by western blot (C) and cells were analyzed by flow cytometry for the percentage of GFP-positive cells (D). (C) The immunoblot shown is representative of at least 3 independent VLP productions. (bottom) Quantification of TASOR degradation rate from 3 experiments. A one-sample t-test was performed. Data are presented as mean ± standard deviation (SD). Statistical significance is indicated as follows: *p < 0.05, **p < 0.01, ***p < 0.001. (D) The reactivation rate corresponds to the percentage of GFP-positive cells in the presence of one viral protein over the percentage obtained without viral protein (empty condition). Reactivation rates from 3 independent experiments are shown (left), along with raw reactivation data from a representative experiment (right). A one-sample t-test was performed, with statistical significance denoted as: *p < 0.05, **p < 0.01, ***p < 0.001. (E) Indicated HA-Vpr constructs were expressed in HeLa cells, then an anti-HA immunoprecipitation was performed, and proteins were revealed by western-blot.

Overall, recurrent adaptation cycles to antagonize restriction factors during lentiviral evolution have selected different determinants (summarized in Fig. 5D).

We next explored the specificity and determinants against SAMHD1 and TASOR within the HIV-2 clade, which is a single lentiviral lineage as opposed to SIVagms. Specifically, we tested whether Vpx from different HIV-2 strains could have different abilities to induce TASOR degradation. We tested a panel of seven HIV-2s from the A or B pandemic groups (sequences Sup Fig. 1): control Vpx from Ghana-1 (group A), Rod (group A) and JK (group B) strains and four Vpx alleles of HIV-2 derived from PLWH-2, which were selected from a study by Yu *et al* as able to induce the degradation of SAMHD1 (30). Two alleles were from effective controllers (EC: high CD4+T cells and undetectable viral load for several years) and two from non-controllers (NC: low CD4 count and high viral load) (30). Vpx from Ghana-1, Rod and JK strains, one NC Vpx (RH2-1-D8, group A) and one EC Vpx (RH2-135C1, group A) were able to induce TASOR degradation in human J-Lat A1 cells after VLP delivery and subsequently to reactivate HIV-1 in the J-Lat A1 model (Fig. 7). One NC Vpx (RH2-7B3, group A) and one EC Vpx (RH2-22-1B4, group B) were impaired in their ability to induce TASOR degradation and reactivation of the latent virus (Fig. 7, strongly impaired for B3, poorly for B4). Therefore, the efficiency of Vpx-mediated TASOR degradation and HIV reactivation was variable within the HIV-2 lineage and did not correlate with the potency of viral control in PLWH-2. We further took advantage of these phenotypic differences and constructed chimera to identify Vpx determinants. In particular, we used NC Vpx RH2-1-D (competent for TASOR degradation) and NC Vpx RH2-7B3 (non competent). The two share 84.8 % identity, with most differences (highlighted in bold in Fig. 8A) in two regions at the N-ter (N-ter tail + first part of helix-1) and the C-ter (helix 3 + C-ter tail). We found that the replacement of the N-ter region (chimera D8B3), but not the C-ter (chimera B3D8B3), from Vpx RH2-7B3 by RH2-1-D8 led to TASOR degradation and reactivation of latent HIV-1 (gain-of-function, Fig. 8; C: incorporation, D: degradation, E: reactivation), showing that RH2-1-D8 Vpx N-ter has key determinants in TASOR antagonism. Yet, in the context of another chimera (B4D8B4, Fig. 9), we revealed that the C-ter region was this time important for improving TASOR degradation efficiency (Figure 9). Overall, the capacity to trigger TASOR degradation is context dependent and relies on N-terminal and C-terminal regions of the viral protein.

**Figure 7:**
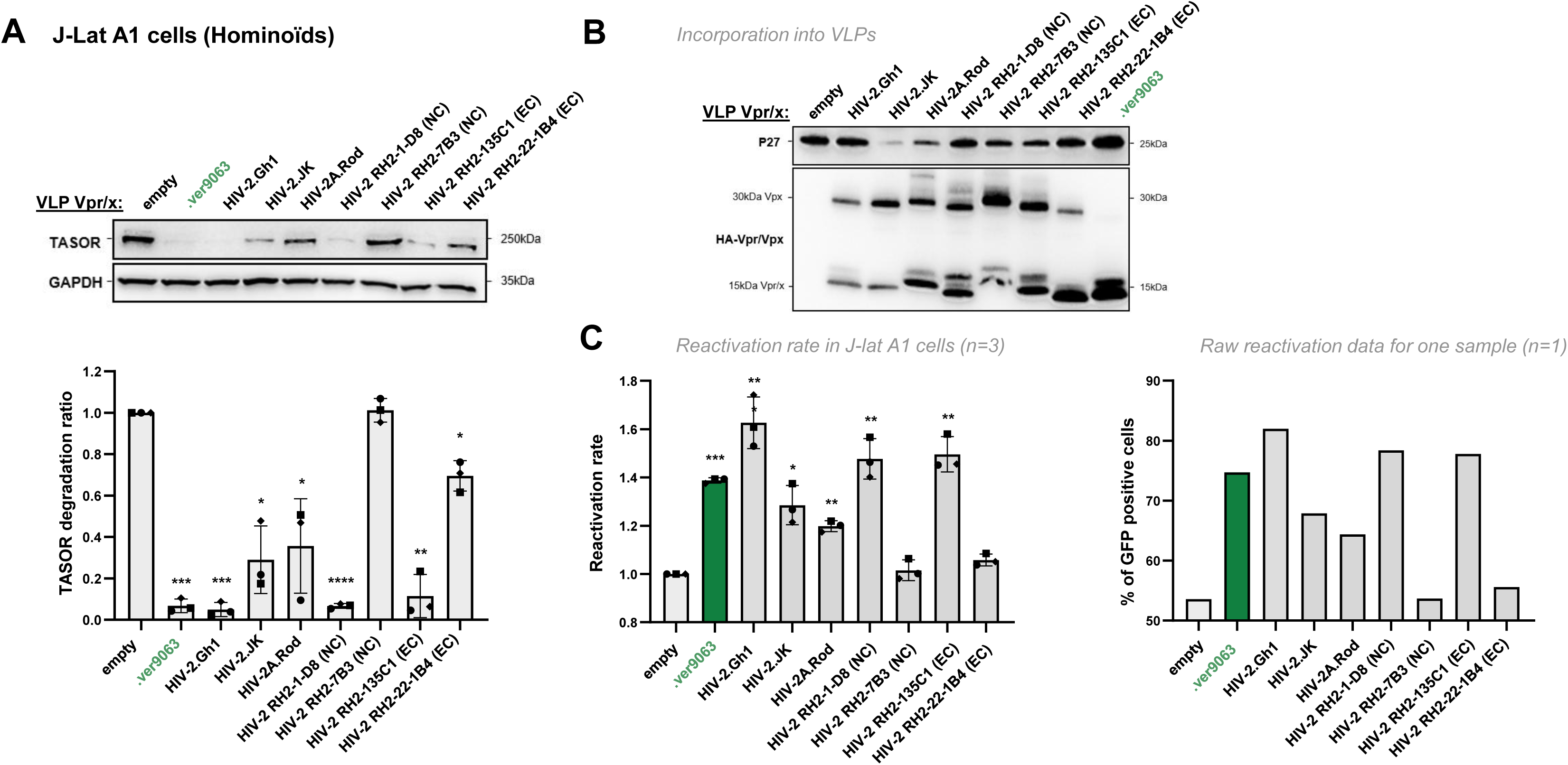
TASOR degradation by Vpx proteins from different origins, including HIV-2 from PLWH-2. Human J-Lat A1 T cells were treated with VLPs containing Vpx proteins from different origins (control Vpx from Ghana-1, Rod and JK strains and four HIV-2 Vpx alleles derived from PLWH-2, according to Yu *et al.*) and overnight with TNF-α. Whole-cell extracts and Vpx incorporation into VLPs were analyzed by western blot (A and B) and cells were also analyzed by flow cytometry for the percentage of GFP-positive cells (C). (A) (Top) the immunoblot shown is representative of at least 3 independent experiments; (bottom) Quantification of TASOR degradation rate from 3 experiments. A one-sample t-test was performed. Data are presented as mean ± standard deviation (SD). Statistical significance is indicated as follows: *p < 0.05, **p < 0.01, ***p < 0.001. (B) Incorporation of viral proteins into the VLPs used in 9A is checked by western-blot. (C) The reactivation rate corresponds to the percentage of GFP-positive cells in the presence of one viral protein over the percentage obtained without viral protein (empty condition). Reactivation rates from 3 independent experiments are shown (left), along with raw reactivation data from one representative experiment (right).

**Figure 8:**
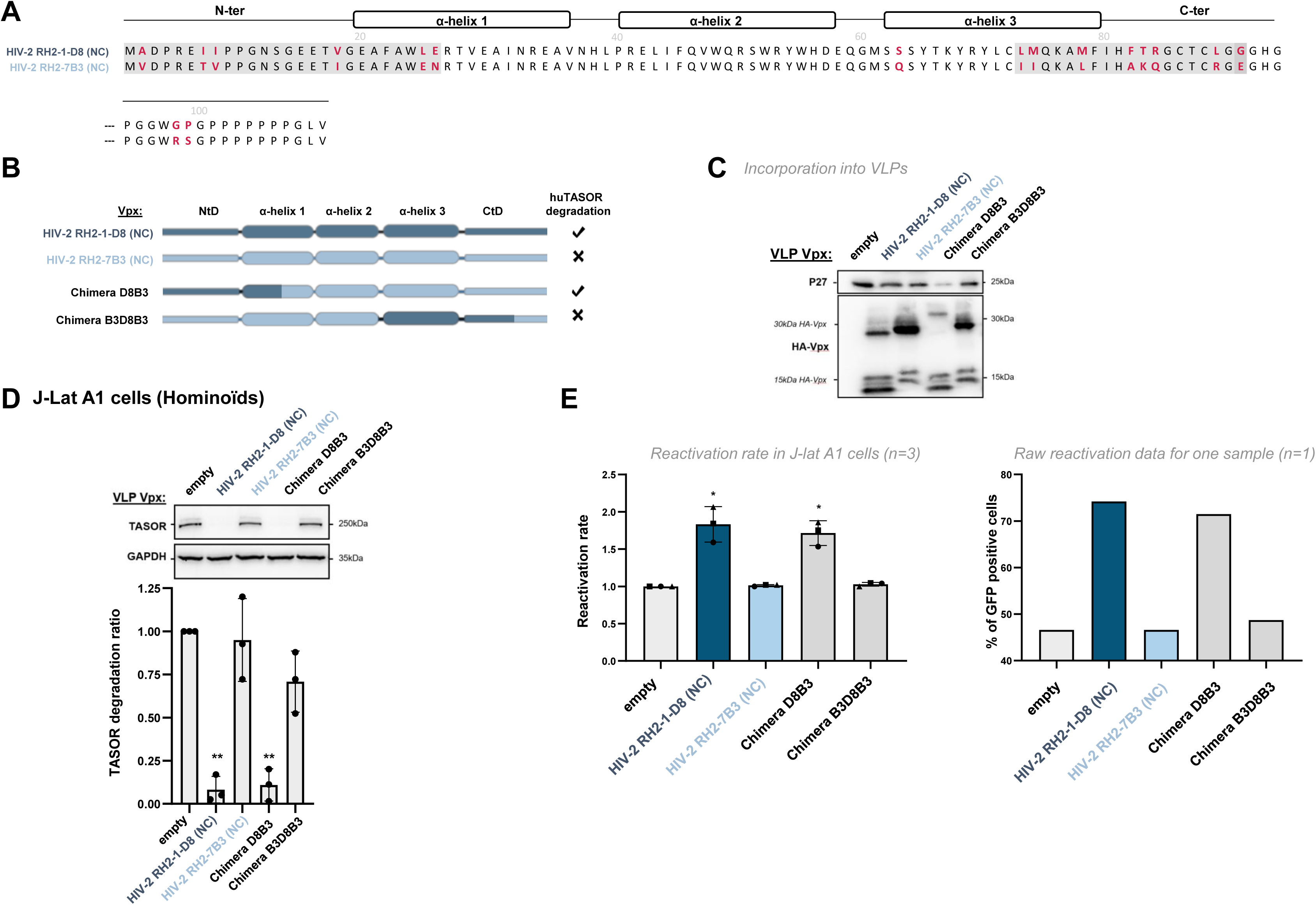
The N-ter region of Vpx RH2-1D8 confers on Vpx RH2-7B3 the ability to induce the degradation of the HUSH core protein TASOR. (A) Sequences from Vpx RH2-1D8 and Vpx RH2-7B3 were aligned to point out amino acid differences (in red). Cross-referenced sequences for chimera are highlighted in light gray. (B) Representation of the two indicated Vpx proteins and chimera. (C, D, E) Vpx incorporation into VLPs, TASOR degradation in J-Lat A1 T cells following Vpx delivery by VLPs and HIV-1 reactivation were assessed as in Figure 7.

**Figure 9:**
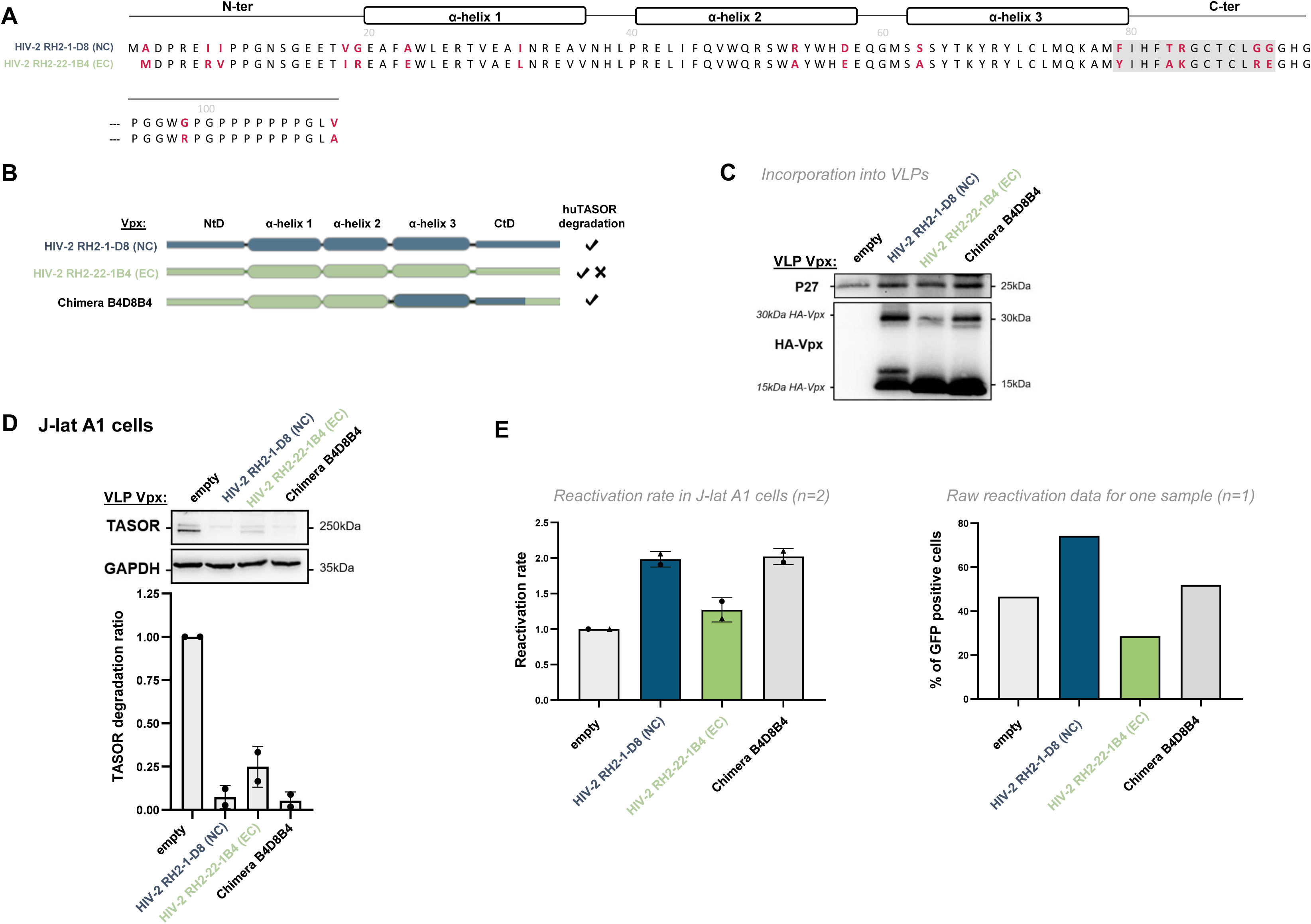
The C-ter region of Vpx RH2-1D8 confers on Vpx RH2-22-1B4 the ability to induce the degradation of the HUSH core protein TASOR. (A) Sequences from Vpx RH2-1D8 and Vpx RH2-22-1B4 were aligned to point out amino acid differences (in red). The cross-referenced sequence for chimera is highlighted in light gray. (B) Representation of the two indicated Vpx proteins and chimera. (C, D, E) Vpx incorporation into VLPs, TASOR degradation in J-Lat A1 T cells following Vpx delivery by VLPs and HIV-1 reactivation were assessed as in Figure 7.

This study and previous work (27, 28) showed differences between Vpr/Vpx proteins from various lentiviral lineages to counteract human TASOR (“lentiviral lineage-specificity”). To complete the virus-host heterologous assays, we next tested the ability of lentiviral SIVagm Vpr proteins to degrade TASOR from divergent primate host species, using cells from New World monkeys (NWMs; owl monkey *Aotus trivirgatus* (aotTri) kidney OMK cells) in addition to cells used up to now from hominoids (human J-Lat A1 cells) and Old World monkeys (OWMs; AGM Vervet Vero cells). Of note, in OMK cells, cyclosporin A (CsA) was added before the delivery of lentiviral proteins to bypass the Trim-CypA block responsible for capsid destabilization upon entry (36). First, we found that SIVagm Vpr proteins displayed the same phenotypes in OMK cells as in VERO and J-Lat A1 cells in presence of CsA (Fig. 10A: incorporation, Fig. 10B: degradation in OMK cells). Therefore, we extended our panel of lentiviral proteins by testing Vpx proteins from the HIV-2/SIVsmm lineage (HIV-2 Ghana-1 strain and SIVsmm Vpx). As previously shown by us and others (27, 28, 37), HIV-2 and SIVsmm Vpx induced human TASOR degradation (Fig. 10C); in addition, we show here that these viral proteins present the same phenotypes in AGM Vervet cells (VERO) (Fig. 10D). Moreover, although SIVagm.Ver, SIVagm.Tan and SIVagm.Sab Vpr proteins had similar phenotypes in owl monkey, AGM and human cells, Vpx proteins from HIV-2 and SIVsmm were unable to induce TASOR degradation in owl monkey cells as opposed to in human and AGM Vervet cells (Fig. 10E). To further confirm TASOR degradation phenotypes in owl monkey cells, we incorporated SIVagm.Ver Vpr and HIV-2 Vpx in SIV-derived GFP encoding viruses and found that GFP expression in owl monkey cells was similar (Fig. 10F). This ruled out the hypothesis that Vpx delivery was impaired in owl monkey cells, though we cannot exclude the possibility that the stability of the two viral proteins may differ.

**Figure 10:**
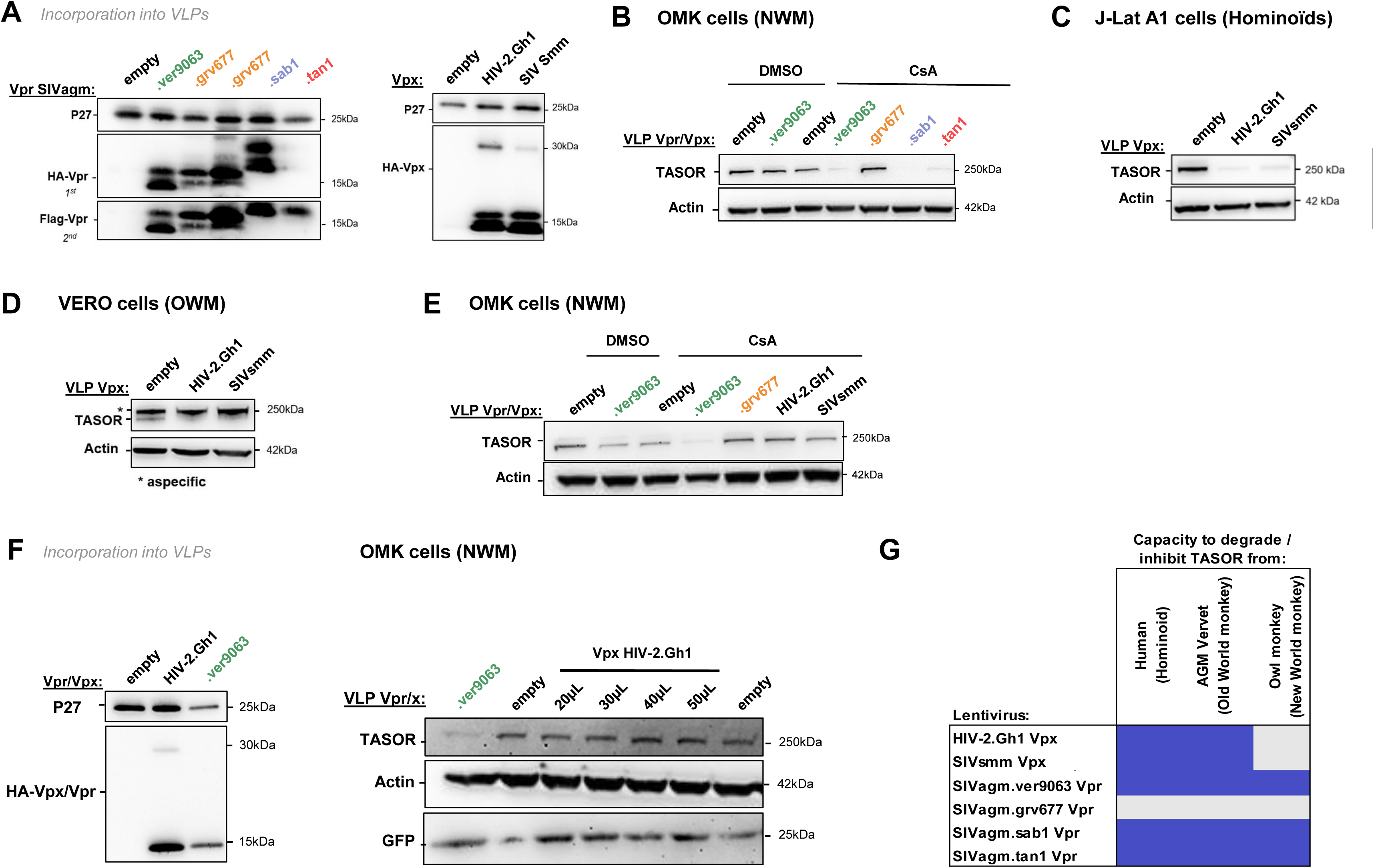
HIV-2/SIVsmm Vpx induces TASOR degradation in human and VERO cells, but not in Owl monkey cells. (A) Incorporation of Vpr and Vpx proteins into VLPs as described in Figure 3A. (B) OMK cells were treated with VLPs containing the indicated Vpr/Vpx proteins. After overnight treatment with cyclosporine CsA (or DMSO in control samples), whole-cell extracts were analyzed by western blot. (C) Human J-Lat A1 T cells were treated with VLPs containing Vpx proteins. After overnight treatment with TNF-α, whole-cell extracts were analyzed by western blot, the immunoblot shown is representative of at least 3 independent VLP productions. (D) Same as (C), but in VERO cells. (E) same as B, but with a different set of viral proteins. (F) (Left) SIVagm.Ver Vpr or HIV-2 Vpx were incorporated into SIV-derived viruses that express GFP following cell transduction. Vpx or Vpr incorporation was checked by western blot on the VLPs. (Right) OMK cells were treated with the Vpx or Vpr-containing SIV-derived viruses. After overnight treatment with CsA, whole-cell extracts were analyzed by western blot. (G) Summary of the degradation phenotypes of the lentiviral proteins. Blue: efficient degradation of TASOR and gray: no degradation.

Altogether, on top of lentivirus-lineage specificity, we now have one evidence of host-species specificity in the interplay between the HUSH complex and the lentiviral Vpr/Vpx proteins (summary Fig. 10G).

Next, we set out to determine whether the apparent host-species specificity could result from a defect of binding of Vpx to DCAF1 and/or TASOR. First, both SIVagm.Ver Vpr and HIV-2 Vpx could interact with owl monkey DCAF1 (Fig. 11A). Unfortunately, interaction of viral proteins with endogenous TASOR is difficult to detect. Thus, we generated by gene synthesis a vector encoding a flag-tagged owl monkey TASOR. The TASOR sequence of the *Aotus trivirgatus* species could not be obtained, we used the publicly-available sequence from a closely related owl monkey species, *Aotus nancymaae* (aotNan; XM_012460781.1). We conducted experiments involving the simultaneous exogenous expression of Flag-tagged owl monkey (aotNan) or human TASOR with, or without, the HA-tagged HIV-2 Vpx or SIVagm.Ver Vpr. By an anti-Flag co-immunoprecipitation assay, we found that both HIV-2 Vpx and SIVagm.Ver Vpr exhibited interaction with human Flag-TASOR but also with owl monkey (aotNan) Flag-TASOR (Fig. 11B). These findings suggest that HIV-2 Vpx is able to interact with owl monkey DCAF1 and (aotNan) TASOR despite the lack of degradation of TASOR in owl monkey (aotTri) cells.

**Figure 11:**
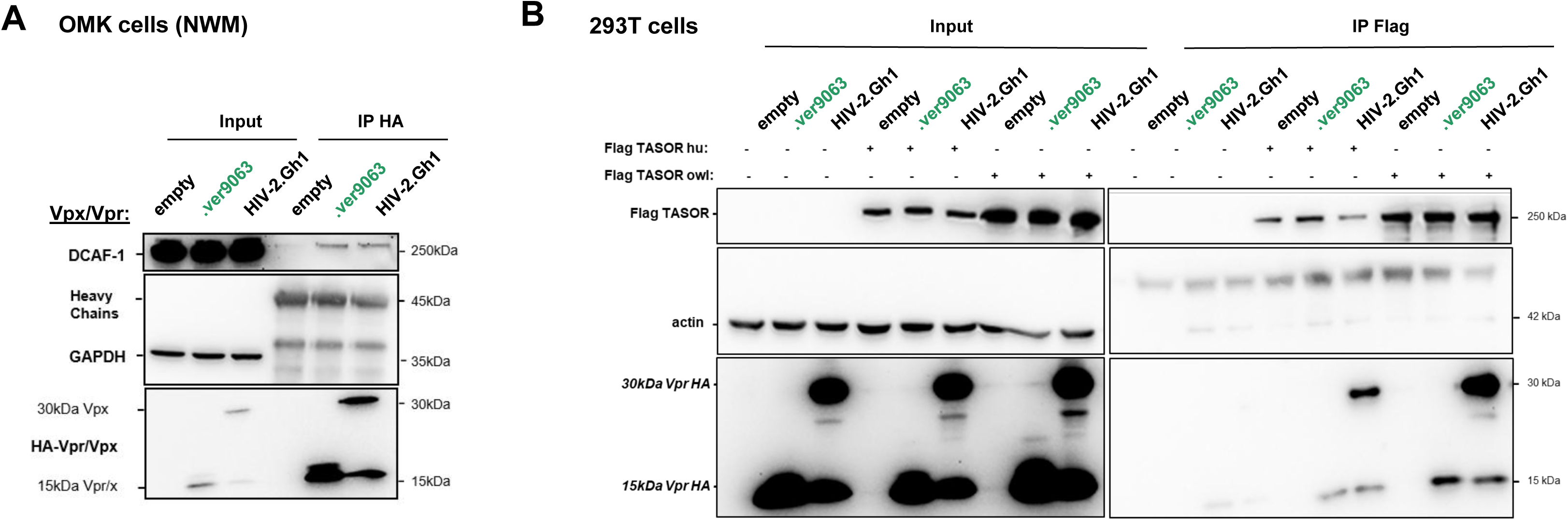
HIV-2 Vpx interacts with TASOR and DCAF1 in Owl monkey cells. (A) Indicated HA-Vpr/Vpx constructs were expressed in OMK cells, then an anti-HA immunoprecipitation was performed, and proteins were revealed by western-blot. (B) SIVagm.Ver9063 Vpr or HIV-2.Gh1 Vpx were overexpressed in OMK cells with human or owl Flag-TASOR, then an anti-Flag immunoprecipitation was performed, and proteins were revealed by western-blot.

## DISCUSSION

In the molecular arms-race between pathogenic viruses and their hosts, proteins are submitted to strong selective pressures. Adaptive mutations in the host immune defense may result from the escape from a viral antagonist, while adaptive changes in viral proteins may maintain the virus’ ability to counteract the host defense. Taking advantage of natural variations in lentiviruses and host immune defenses from African green monkeys at the inter-and intra-species levels, we show that closely related Vpr proteins can induce the degradation of SAMHD1 variants (haplotypes IV and V) through distinct molecular determinants and that the SIVagm.Ver Vpr protein uses also distinct determinants to trigger SAMHD1 and TASOR degradation. Lastly, we discovered that HIV-2 Vpx proteins from people living with HIV present distinct TASOR degradation abilities, which do not correlate with viremia. TASOR-antagonism determinants are context dependent and are both in the N and C termini. Overall, our results underline the high plasticity of Vpr and Vpx proteins to hijack ubiquitin ligase complexes and to eliminate restrictive host proteins. Lastly, we have evidence of host species-specificity in HUSH antagonism with HIV-2 and SIVsmm Vpx proteins able to induce TASOR degradation in human and AGM VERO cells, but not in owl monkey OMK cells.

Only four amino acid differences specify the distinct behavior of the SAMHD1 haplotypes IV and V, with respect to SIVagm Vpr proteins (26). We found that SIVagm.Ver and SIVagm.Tan Vpr both use their C-ter tails to target SAMHD1 haplotype IV. Interestingly, in the crystal structures of the ternary complexes of DCAF1, SAMHD1 and Vpx from SIVsmm, which also targets the C-ter domain of SAMHD1, no residue of the C-terminal tail (also VR3) of Vpx were found in contact with SAMHD1 (23, 24). Nonetheless, such structural studies might not be adapted to Vpx C-ter tail, due to its flexible nature. It is also possible that the recognition mechanism has diverged within the SIVagm lineage or that the C-terminal tail has only a structural role in maintaining the recognition interface.

Our approach using chimeric proteins did not allow to reveal specific viral determinants of SIVagm.Gri Vpr involved in haplotype V degradation, suggesting that the integrity of different parts of the viral protein is required. The use of different interfaces by closely related viral proteins underlines how antagonism of a given restriction factor results from different modes of adaptation by the virus. Our results showing that the substitution of only one amino-acid within a given Vpr protein restores its ability to induce the degradation of SAMHD1 further supports this model of molecular adaptation along evolution.

While the C-terminal tail of SIVagm.Ver Vpr conferred on the SIVagm.Gri protein the ability to induce SAMHD1 haplotype IV degradation, the α-helix 3 of SIVagm.Ver Vpr conferred on the SIVagm.Gri protein the ability to degrade the HUSH core protein TASOR. These results suggest that the C-terminal tail and α-helix 3 are key determinants for SAMHD1 haplotype IV and TASOR degradation, respectively, but they do not exclude the possibility that the integrity of other determinants within the viral protein is required for the degradation of each substrate. Indeed, as the SIVagm.Ver and SIVagm.Gri proteins are very similar in sequence, chimera may share similar determinants important for the viral protein activity. In any event, the use of distinct determinants for the degradation of two different substrates fits with a model of ubiquitin ligase hijacking, in which a viral protein would induce the degradation of different host factors using distinct viral interfaces for substrate recognition, but hijacking only one type of ubiquitin ligase. Intriguingly, SIVagm.Gri Vpr could not induce HUSH degradation, while the integrity of the whole protein seemed required for SAMHD1 haplotype V degradation. Therefore, the selective pressure imposed on the “entire” virus protein to counteract SAMHD1 may have limited its ability to adapt and counteract HUSH (27). Alternatively, it is possible that SIVagm.Gri Vpr cannot degrade human, Vervet AGM or owl monkey TASORs, but can degrade the TASOR from Grivet AGMs, in a host-species specific manner. To address this question, all three components of HUSH (TASOR, MPP8 and periphilin) should be analyzed for polymorphisms and variants within and between AGM species and more extensively in primates, and tested for their degradation in the presence of the different SIVagm Vpr proteins. It is also possible that important cofactors are lacking/different in the cells from other hosts. Altogether, the use of different viral interfaces between closely related Vpr proteins within the SIVagm lineage highlights the dynamism and constraints of the molecular interactions between Vpr proteins, SAMHD1 and HUSH, as a result of a cat-and-mouse game during evolution.

We further highlighted that Vpx proteins of HIV-2 from PLWH-2 harbor different phenotypes. The ability to induce TASOR degradation can be restored after substitution of residues either in the N or in the C-ter part (including helix 3) of the viral protein depending on the Vpx, suggesting that different regions in Vpx could contribute to its interaction with HUSH.

Our study also opens questions concerning the role of HIV-2 Vpx. By inducing HUSH degradation, Vpx is able to increase the proviral transcription from the HIV LTR and the stability of the LTR-driven transcript, thus Vpx counteracts the silencing of the provirus (38). One could think at first that Vpx confers an advantage in HIV-2 replication by enhancing LTR-driven RNA expression. However, interestingly, this study of Vpx proteins derived from both viremic (NCs) or long-term aviremic (EV) PLWH-2 showed that the capacity to induce TASOR degradation did not correlate with the viremia. The importance of TASOR degradation over the course of HIV-2 infections remains to be investigated. In addition, we cannot exclude that some HIV-2 strains may use other non-Vpx determinants in TASOR antagonism.

Lastly, we determined here that HUSH antagonism presents some features of host-species specificity in that the HIV-2/SIVsmm Vpx proteins could induce the degradation of human and AGM vervet, but not of owl monkey TASOR. The underlying causes remain unknown. It is unlikely a defect in the viral protein delivery, because other Vpr/Vpx proteins were well delivered and functional in these cells, but we cannot exclude that HIV-2/SIVsmm Vpx may be particularly unstable in OMK cells. Our coIP results suggest that absence of degradation does not result from a lack of interaction with DCAF1 or TASOR (35). However, one limitation in our coIP experiments is that, due to challenges in sequencing the entire OMK TASOR, we here tested TASOR from a publicly-available sequence of *Aotus nancymaae* TASOR, which could be slightly different from that of OMK cells (*Aotus trivirgatus*).

Virus-host species-specificity is a hallmark of restriction factors. Antagonism of APOBEC3G by Vif occurs in a species-specific manner in Catharine primates and AGM populations, and functional evolutionary studies showed that the specificity of Vif reflects adaptation of the virus to the host including in key cross-species transmission events (3, 39–46). Experimental virus-host heterologous *in vivo* infection of AGMs showed that the lentiviral *vif* gene can adapt to Vif-resistant APOBEC3G haplotypes (43). Although not observed in the same experimental setting with Vpr (26), SIVagm Vpr proteins have acquired distinct interfaces to counteract SAMHD1, suggesting some adaptation of the virus to SAMHD1 selective pressure. The resistance of owl monkey TASOR to HIV-2/SIVsmm Vpx further suggests that HUSH may have evolved in some primate ancestors in response to past lentiviral epidemics. Alternatively, another pathogen, endogenous viral elements or drivers may have shaped primate TASOR resulting in some host species-specificity. More primate sequences in certain lineages and evolutionary analyses combined with functional assays would help to determine the modes and causes of such host evolution. Altogether, on one hand, SAMHD1 and HUSH antagonisms by Vpr/Vpx proteins appear to be critical components of primate lentiviral fitness witnessed by the complex dynamism of the interactions at stake. On the other hand, results from this study and others (30) suggest that Vpx-mediated SAMHD1 or HUSH antagonism does not always correlate with viral control in PLWH-2. Future work will help to resolve this apparent paradox.

## ACKNOWLEDGMENTS

We thank all the members of the “Retrovirus, Infection and Latency team” for fruitful discussions. We also thank the members of the LP2L team at CIRI, Lyon, for support. We acknowledge the CYBIO, IMAG’IC core facilities of the Institut Cochin.

This work was supported by grants from the SIDACTION, the French Research Agency on HIV and Emerging Infectious Diseases ANRS/MIE and Fondation pour la Recherche Médicale (FRM, EQU202203014684 attributed to F.MG.). P.L. was supported by Université de Paris Cité, M.M.M. by SIDACTION, C.G. by FRM (EQU202203014684 attributed to F.MG.), K.Z. by ANRS/MIE, R.M. by SIDACTION and FRM (EQU202203014684 attributed to F.MG.). The work in the laboratory of L.E. is supported by grants from the ANRS/MIE (#ECTZ118944 to LE) and SIDACTION (n°21-1-AEQ-12972-2 to LE and FMG).

---

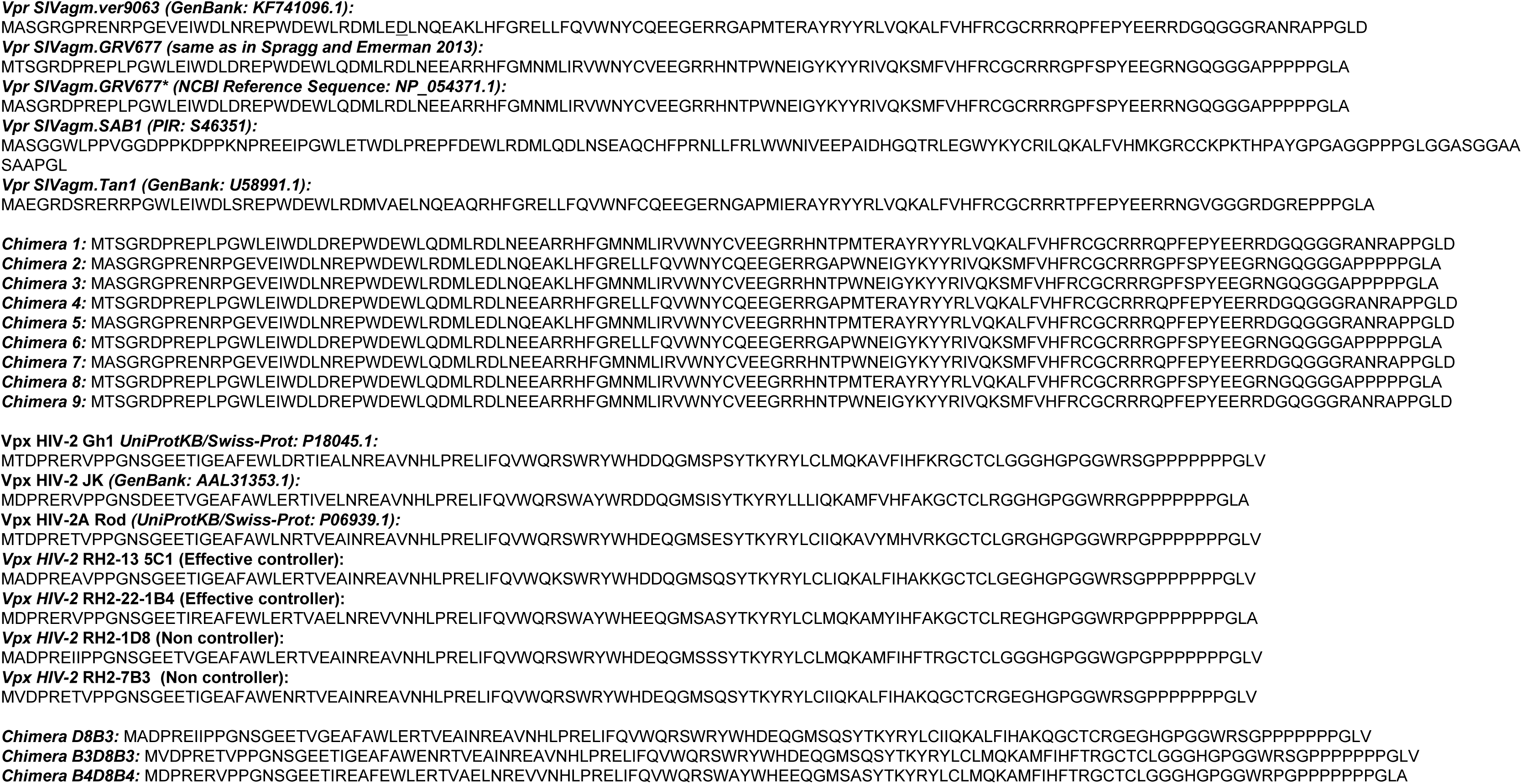

## Notes

### Competing Interest Statement

The authors have declared no competing interest.

### Summary of Updates

In the first version, we were characterizing viral determinants involved in SAMHD1 and HUSH antagonism, using chimeric Vpr proteins between SIVagm.Ver and SIVagm.Gri lentiviruses. We have now quantified most of the western-blots and performed statistical analyses. We also have checked that the chimera were all able to bind DCAF1. Importantly, we consider that a chimera is functional only when the protein is able to induce the degradation of at least one given substrate. Moreover, new mutations have been introduced in the parental Vpr proteins. One of the main addition is the study of Vpx proteins of HIV-2 from people living with HIV-2 (PLWH-2). We found that these Vpx proteins have different abilities to degrade HUSH, which led us to use the same chimera strategy to decipher Vpx determinants required for HUSH degradation. Phenotypes rely on small changes in either the N or C terminal part of Vpx, depending on the context. Interestingly, along this study, we discovered that the ability to induce TASOR degradation did not correlate with viremia. In the previous version, it was described that HIV-2 and SIVsmm Vpx degrading HUSH from human and vervet monkey cells cannot not degrade HUSH in owl monkey cells, suggesting some host species-specificity. We now show that SIVagm.Ver Vpr or HIV-2 Vpx interacts with DCAF1 or overexpressed TASOR from owl monkey cells. Thus, we could not point out an explanation for this host-species specificity.

## REFERENCES

1. M. D. Daugherty, H. S. Malik, Rules of engagement: molecular insights from host-virus arms races. Annu Rev Genet 46, 677–700 (2012).

2. N. K. Duggal, M. Emerman, Evolutionary conflicts between viruses and restriction factors shape immunity. Nat Rev Immunol 12, 687–695 (2012).

3. L. Etienne et al., The Role of the Antiviral APOBEC3 Gene Family in Protecting Chimpanzees against Lentiviruses from Monkeys. PLoS Pathog 11, e1005149 (2015).

4. A. Kirmaier et al., TRIM5 suppresses cross-species transmission of a primate immunodeficiency virus and selects for emergence of resistant variants in the new species. PLoS Biol 8 (2010).

5. R. M. Pinto et al., BTN3A3 evasion promotes the zoonotic potential of influenza A viruses. Nature 10.1038/s41586-023-06261-8 (2023).

6. D. Sauter, F. Kirchhoff, Key Viral Adaptations Preceding the AIDS Pandemic. Cell Host Microbe 25, 27–38 (2019).

7. L. Etienne, B. H. Hahn, P. M. Sharp, F. A. Matsen, M. Emerman, Gene loss and adaptation to hominids underlie the ancient origin of HIV-1. Cell Host Microbe 14, 85–92 (2013).

8. V. Planelles et al., Vpr-induced cell cycle arrest is conserved among primate lentiviruses. J Virol 70, 2516–2524 (1996).

9. G. L. Stivahtis, M. A. Soares, M. A. Vodicka, B. H. Hahn, M. Emerman, Conservation and host specificity of Vpr-mediated cell cycle arrest suggest a fundamental role in primate lentivirus evolution and biology. J Virol 71, 4331–4338 (1997).

10. Y. Zhu et al., Comparison of cell cycle arrest, transactivation, and apoptosis induced by the simian immunodeficiency virus SIVagm and human immunodeficiency virus type 1 vpr genes. J Virol 75, 3791–3801 (2001).

11. V. M. Hirsch et al., Vpx is required for dissemination and pathogenesis of SIV(SM) PBj: evidence of macrophage-dependent viral amplification. Nat Med 4, 1401–1408 (1998).

12. T. Schaller, H. Bauby, S. Hue, M. H. Malim, C. Goujon, New insights into an X-traordinary viral protein. Front Microbiol 5, 126 (2014).

13. K. Hrecka et al., Vpx relieves inhibition of HIV-1 infection of macrophages mediated by the SAMHD1 protein. Nature 474, 658–661 (2011).

14. N. Laguette et al., SAMHD1 is the dendritic-and myeloid-cell-specific HIV-1 restriction factor counteracted by Vpx. Nature 474, 654–657 (2011).

15. E. S. Lim et al., The ability of primate lentiviruses to degrade the monocyte restriction factor SAMHD1 preceded the birth of the viral accessory protein Vpx. Cell Host Microbe 11, 194–204 (2012).

16. H. M. Baldauf et al., SAMHD1 restricts HIV-1 infection in resting CD4(+) T cells. Nat Med 18, 1682–1689 (2012).

17. H. Lahouassa et al., SAMHD1 restricts the replication of human immunodeficiency virus type 1 by depleting the intracellular pool of deoxynucleoside triphosphates. Nat Immunol 13, 223–228 (2012).

18. A. Bergamaschi et al., The human immunodeficiency virus type 2 Vpx protein usurps the CUL4A-DDB1 DCAF1 ubiquitin ligase to overcome a postentry block in macrophage infection. J Virol 83, 4854–4860 (2009).

19. E. Le Rouzic et al., HIV1 Vpr arrests the cell cycle by recruiting DCAF1/VprBP, a receptor of the Cul4-DDB1 ubiquitin ligase. Cell Cycle 6, 182–188 (2007).

20. J. Ahn et al., HIV/simian immunodeficiency virus (SIV) accessory virulence factor Vpx loads the host cell restriction factor SAMHD1 onto the E3 ubiquitin ligase complex CRL4DCAF1. J Biol Chem 287, 12550–12558 (2012).

21. S. Srivastava et al., Lentiviral Vpx accessory factor targets VprBP/DCAF1 substrate adaptor for cullin 4 E3 ubiquitin ligase to enable macrophage infection. PLoS Pathog 4, e1000059 (2008).

22. O. I. Fregoso et al., Evolutionary toggling of Vpx/Vpr specificity results in divergent recognition of the restriction factor SAMHD1. PLoS Pathog 9, e1003496 (2013).

23. D. Schwefel et al., Molecular determinants for recognition of divergent SAMHD1 proteins by the lentiviral accessory protein Vpx. Cell Host Microbe 17, 489–499 (2015).

24. D. Schwefel et al., Structural basis of lentiviral subversion of a cellular protein degradation pathway. Nature 505, 234–238 (2014).

25. N. Laguette et al., Evolutionary and functional analyses of the interaction between the myeloid restriction factor SAMHD1 and the lentiviral Vpx protein. Cell Host Microbe 11, 205–217 (2012).

26. C. J. Spragg, M. Emerman, Antagonism of SAMHD1 is actively maintained in natural infections of simian immunodeficiency virus. Proc Natl Acad Sci U S A 110, 21136–21141 (2013).

27. G. Chougui et al., HIV-2/SIV viral protein X counteracts HUSH repressor complex. Nat Microbiol 3, 891–897 (2018).

28. L. Yurkovetskiy et al., Primate immunodeficiency virus proteins Vpx and Vpr counteract transcriptional repression of proviruses by the HUSH complex. Nat Microbiol 3, 1354–1361 (2018).

29. I. A. Tchasovnikarova, et al., GENE SILENCING. Epigenetic silencing by the HUSH complex mediates position-effect variegation in human cells. Science 348, 1481–1485 (2015).

30. H. Yu et al., The efficiency of Vpx-mediated SAMHD1 antagonism does not correlate with the potency of viral control in HIV-2-infected individuals. Retrovirology 10, 27 (2013).

31. T. Gramberg, N. Sunseri, N. R. Landau, Evidence for an activation domain at the amino terminus of simian immunodeficiency virus Vpx. J Virol 84, 1387–1396 (2010).

32. P. R. Johnson et al., Simian immunodeficiency viruses from African green monkeys display unusual genetic diversity. J Virol 64, 1086–1092 (1990).

33. C. H. Douse et al., TASOR is a pseudo-PARP that directs HUSH complex assembly and epigenetic transposon control. Nat Commun 11, 4940 (2020).

34. A. Jordan, D. Bisgrove, E. Verdin, HIV reproducibly establishes a latent infection after acute infection of T cells in vitro. EMBO J 22, 1868–1877 (2003).

35. M. M. Martin et al., Binding to DCAF1 distinguishes TASOR and SAMHD1 degradation by HIV-2 Vpx. PLoS Pathog 17, e1009609 (2021).

36. D. M. Sayah, E. Sokolskaja, L. Berthoux, J. Luban, Cyclophilin A retrotransposition into TRIM5 explains owl monkey resistance to HIV-1. Nature 430, 569–573 (2004).

37. E. J. D. Greenwood et al., Promiscuous Targeting of Cellular Proteins by Vpr Drives Systems-Level Proteomic Remodeling in HIV-1 Infection. Cell Rep 27, 1579–1596 e1577 (2019).

38. R. Matkovic et al., TASOR epigenetic repressor cooperates with a CNOT1 RNA degradation pathway to repress HIV. Nat Commun 13, 66 (2022).

39. H. P. Bogerd, B. P. Doehle, H. L. Wiegand, B. R. Cullen, A single amino acid difference in the host APOBEC3G protein controls the primate species specificity of HIV type 1 virion infectivity factor. Proc Natl Acad Sci U S A 101, 3770–3774 (2004).

40. R. Mariani et al., Species-specific exclusion of APOBEC3G from HIV-1 virions by Vif. Cell 114, 21–31 (2003).

41. H. Xu et al., A single amino acid substitution in human APOBEC3G antiretroviral enzyme confers resistance to HIV-1 virion infectivity factor-induced depletion. Proc Natl Acad Sci U S A 101, 5652–5657 (2004).

42. A. A. Compton, M. Emerman, Convergence and divergence in the evolution of the APOBEC3G-Vif interaction reveal ancient origins of simian immunodeficiency viruses. PLoS Pathog 9, e1003135 (2013).

43. A. A. Compton, V. M. Hirsch, M. Emerman, The host restriction factor APOBEC3G and retroviral Vif protein coevolve due to ongoing genetic conflict. Cell Host Microbe 11, 91–98 (2012).

44. M. D’Arc et al., Origin of the HIV-1 group O epidemic in western lowland gorillas. Proc Natl Acad Sci U S A 112, E1343–1352 (2015).

45. M. Letko, T. Booiman, N. Kootstra, V. Simon, M. Ooms, Identification of the HIV-1 Vif and Human APOBEC3G Protein Interface. Cell Rep 13, 1789–1799 (2015).

46. M. Letko et al., Vif proteins from diverse primate lentiviral lineages use the same binding site in APOBEC3G. J Virol 87, 11861–11871 (2013).

